# Implementation of the β-hydroxyaspartate cycle increases growth performance of *Pseudomonas putida* on the PET monomer ethylene glycol

**DOI:** 10.1101/2022.08.08.503134

**Authors:** Lennart Schada von Borzyskowski, Helena Schulz-Mirbach, Mauricio Troncoso Castellanos, Francesca Severi, Paul A. Gomez Coronado, Timo Glatter, Arren Bar-Even, Steffen N. Lindner, Tobias J. Erb

## Abstract

Ethylene glycol (EG) is a promising next generation feedstock for bioprocesses. It is a key component of the ubiquitous plastic polyethylene terephthalate (PET) and other polyester fibers and plastics, used in antifreeze formulations, and can also be generated by electrochemical conversion of syngas, which makes EG a key compound in a circular bioeconomy. The majority of biotechnologically relevant bacteria assimilate EG via the glycerate pathway, a wasteful metabolic route that releases CO_2_ and requires reducing equivalents as well as ATP. In contrast, the recently characterized β-hydroxyaspartate cycle (BHAC) provides a more efficient, carbon-conserving route for C2 assimilation. Here we aimed at overcoming the natural limitations of EG metabolism in the industrially relevant strain *Pseudomonas putida* KT2440 by replacing the native glycerate pathway with the BHAC. We first prototyped the core reaction sequence of the BHAC in Escherichia coli before establishing the complete four-enzyme BHAC in *Pseudomonas putida*. Directed evolution on EG resulted in an improved strain that exhibits 35% faster growth and 20% increased biomass yield compared to a recently reported *P. putida* strain that was evolved to grow on EG via the glycerate pathway. Genome sequencing and proteomics highlight plastic adaptations of the genetic and metabolic networks in response to the introduction of the BHAC into *P. putida* and identify key mutations for its further integration during evolution. Taken together, our study shows that the BHAC can be utilized as ‘plug-and-play’ module for the metabolic engineering of two important microbial platform organisms, paving the way for multiple applications for a more efficient and carbon-conserving upcycling of EG in the future.

## 1. Introduction

Plastics are omnipresent. In 2017, the annual world plastic production reached 350 million tons (PlasticsEurope, 2018). Notably, a significant amount of plastic is not disposed in a safe and sustainable manner. It is estimated that almost 80% of the 6,300 million tons of plastic waste that were generated until 2015 were disposed in landfills and elsewhere in the environment (Geyer et al., 2017). Moreover, an estimated 13 million tons of plastic end up in the oceans every year (Danso et al., 2019). This plastic pollution in the environment has serious consequences for the health of organisms and the stability of ecosystems, requiring novel approaches for a sustainable plastic waste management. Microbial degradation and/or upcycling of plastic waste is a promising solution by which the chemical building blocks of plastics are re-used (or upgraded) for the biotechnological production of value-added compounds (Wierckx et al., 2015; Narancic et al., 2017). While different plastic degradation and assimilation pathways were studied in a variety of bacteria (Tiso et al., 2021b), research on microbial valorization of polyethylene terephthalate (PET) is currently most advanced. Following the discovery of the PET-degrading bacterium *Ideonella sakaiensis* (Yoshida et al., 2016), the physiology and biochemistry of PET degradation has been elucidated during the last years. The PET polymer is hydrolyzed into its oligo- and monomers, terephthalate and ethylene glycol (EG), by the enzymes PETase and MHETase, which have been extensively characterized and further engineered for biotechnological applications (Han et al., 2017; Palm et al., 2019; Knott et al., 2020; Yoshida et al., 2021). Yet, for an efficient assimilation and/or upcycling of PET, understanding and engineering the downstream pathways that further convert terephthalate and EG are equally important.

Notably, EG is not only a key building block of PET, as well as other polyester resins and fibers (Yue et al., 2012), but also finds applications as antifreeze agent or solvent worldwide (Dobson, 2000). Furthermore, EG can be electrochemically generated from syngas in a two-step process, thus gaining increasing attention as a key component for a carbon neutral bio-economy (Zheng et al., 2022). Therefore, efforts to develop microbial strains with improved conversion capacities for EG is not only important for microbial PET upcycling, but also in the broader context of establishing circular economic routes for the biotechnological use of this abundant chemical.

Microbial metabolism of EG requires the stepwise oxidation of the C2 compound to glycolaldehyde, glycolate, and ultimately glyoxylate (Figure 1a). These conversions are catalyzed by alcohol and aldehyde dehydrogenases, which employ different cofactors such as NAD^+^, pyrroloquinoline quinone (PQQ), or cytochromes that funnel the electrons into the electron transport chain (ETC). The oxidation of EG has been studied in the Gammaproteobacterium *Pseudomonas putida* KT2440 (Muckschel et al., 2012; Wehrmann et al., 2017; Li et al., 2019), a metabolically versatile bacterium that is frequently used in biotechnology (Belda et al., 2016; Nikel et al., 2018; Weimer et al., 2020). Based on these results, *P. putida* KT2440 has been developed as a microbial chassis organism for EG- and PET-upcycling (Wierckx et al., 2015; Tiso et al., 2021b).

**Figure 1:**
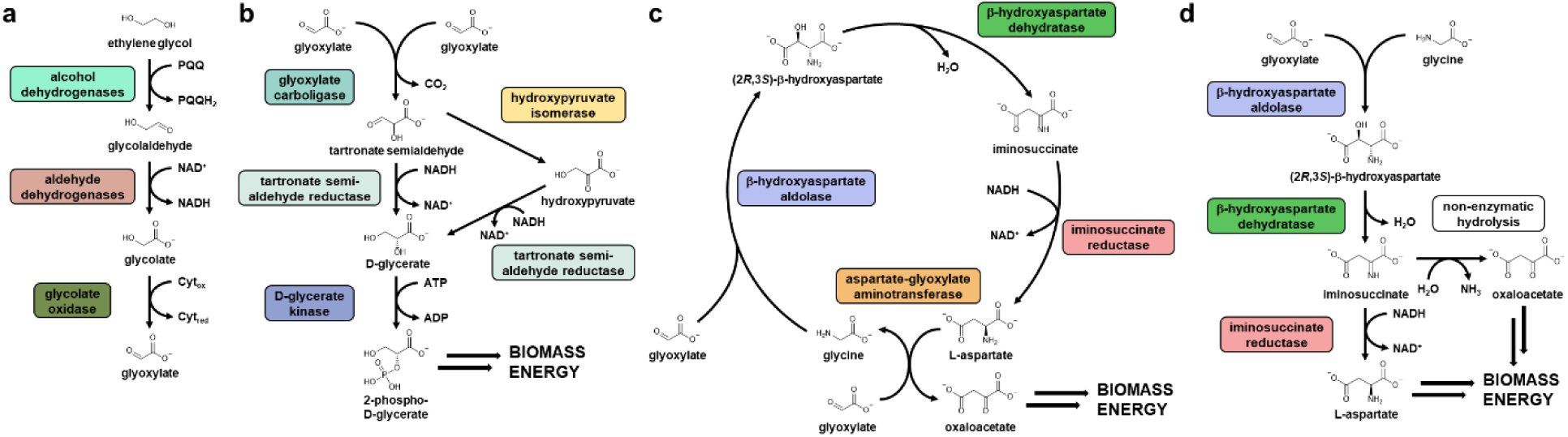
Metabolic pathways involved in EG assimilation. **a**, Oxidative reactions from EG to glyoxylate. Several alcohol and aldehyde dehydrogenases were demonstrated to be involved in the pathway in *P. putida* KT2440 (Muckschel et al., 2012; Wehrmann et al., 2017; Li et al., 2019). **b**, The glycerate pathway, which converts two molecules of glyoxylate into 2-phosphoglycerate and CO_2_. **c**, The BHAC, a cyclic pathway in which four enzymes convert two molecules of glyoxylate into one molecule of oxaloacetate. **d**, The BHA shunt, consisting of three of the BHAC enzymes, is a linear pathway that converts glyoxylate and glycine into aspartate. If non-enzymatic hydrolysis of the labile compound iminosuccinate takes place in the absence of iminosuccinate reductase, oxaloacetate is formed instead of aspartate.

Glyoxylate is further assimilated in *P. putida* KT2440 via the glycerate pathway (Figure 1b) (Muckschel et al., 2012), which is regulated by the transcription factor GclR (Li et al., 2019). EG assimilation in *P. putida* KT2440 has been recently improved through overexpression of the operons encoding for glycerate pathway enzymes and glycolate oxidase (Franden et al., 2018). However, EG assimilation in *P. putida* still suffers from the very first step in the glycerate pathway, the decarboxylating condensation of two molecules of glyoxylate by the enzyme glyoxylate carboligase (Gcl) into tartronate semialdehyde (Gupta et al., 1964; Kaplun et al., 2008). Through this first step, CO_2_ is released, which lowers carbon efficiency of EG assimilation and wastes reducing equivalents. Therefore, we reasoned that replacing the glycerate pathway with a more efficient glyoxylate assimilation pathway would result in improved EG conversion in *P. putida* KT2440.

The β-hydroxyaspartate cycle (BHAC; Figure 1c) is a glyoxylate assimilation module that was partly described in the 1960s (Kornberg et al., 1963; Kornberg et al., 1965) and recently fully characterized in the Alphaproteobacterium *Paracoccus denitrificans*. The pathway employs four enzymes to convert two molecules of glyoxylate into oxaloacetate (Schada von Borzyskowski et al., 2019). Notably, the BHAC does not release CO_2_ and requires only one molecule of NADH to produce oxaloacetate. Moreover, the enzymes of the BHAC were successfully implemented in *Arabidopsis thaliana* recently, demonstrating that this pathway can be transplanted into other organisms (Roell et al., 2021). Altogether, these features make the BHAC a promising alternative for EG assimilation in *P. putida* KT2440.

In this work, we implemented the BHAC in *P. putida* KT2440 to improve biomass yield and growth rate of the organism by 20% and 35% on EG, respectively. We first prototyped the core reaction sequence of the BHAC in different *E. coli* selection strains, before moving the pathway into *P. putida* KT2440. We then used directed evolution to improve BHAC-mediated growth of *P. putida* KT2440 on EG. Finally, we characterized the evolved strain using whole genome sequencing and proteomics to identify the molecular basis for its improved growth behavior. Altogether, our study reports an engineered *P. putida* KT2440 strain that overcomes the limitations of natural EG metabolism through successful implementation of an alternative glyoxylate assimilation module that significantly increases biomass yield and growth rate, laying the foundation for the development of highly efficient chassis for different biotechnological applications in PET and EG conversions.

## 2. Methods

### 2.1. Methods for *E. coli* work (carried out at Max Planck Institutes in Golm and Marburg)

#### 2.1.1. *E. coli* strain construction

All strains used in this study are listed in Supplementary Table 1. Strain SIJ488, which is carrying inducible recombinase and flippase genes (Jensen et al., 2015), was used as wildtype. Gene deletions were performed by λ-Red recombineering or P1 transduction.

#### 2.1.2. Gene deletion by λ-Red recombineering

To delete the genes *aspA, ppc, pck, mdh, mqo,* and *aspC* by recombineering, PCR with pKD3 (pKD3 was a gift from Barry L. Wanner; Addgene plasmid #45604, https://www.addgene.org/45604/) as template, KO primers with 50 bp homologous overhangs and PrimeStar GXL polymerase (Takara Bio) was performed to generate chloramphenicol (CapR) resistance cassettes. To prepare cells for gene deletion, fresh cultures were inoculated in LB, followed by addition of 15 mM L-arabinose at OD ∼0.4-0.5 for induction of the recombinase genes and incubation for 45 min at 37°C. The cells were harvested by centrifugation (11,000 rpm, 30 sec, 2°C) and washed three times with ice cold 10 % glycerol. Electroporation was done with ∼300 ng of Cap cassette PCR-product (1 mm cuvette, 1.8 kV, 25 µF, 200 Ω). Colonies with successful gene deletion were selected for by plating on chloramphenicol-containing plates. Gene deletion was confirmed via one PCR using ‘KO-Ver’ primers (Supplementary Table 2) and one PCR using internal primers (Supplementary Table 2), both with DreamTaq polymerase (Thermo Scientific, Dreieich, Germany). The CapR cassette was removed by addition of 50 mM L-rhamnose to 2 ml LB culture at OD 0.5 for induction of flippase gene expression, followed by incubation for ≥ 3 h at 30 °C. After screening colonies for chloramphenicol sensitivity, successful removal of the CapR cassette was confirmed by PCR using ‘KO-Ver’ primers and DreamTaq polymerase (Thermo Scientific, Dreieich, Germany).

#### 2.1.3. Gene deletion via P1 transduction

*tyrB* and *gcl* were deleted by P1 phage transduction (Thomason et al., 2007). The lysate was produced using strain JW4014 (Δ*tyrB*) or strain JW0495 (Δ*gcl*) from the Keio collection (Baba et al., 2006) with a kanamycin-resistance gene (KmR) in the respective locus. Colonies with the desired deletion were selected for by plating on kanamycin-containing plates. To verify successful gene deletion, the size of the genomic locus was verified by PCR with DreamTaq polymerase (Thermo Scientific, Dreieich, Germany) and the respective KO-Ver primers (Supplementary Table 2). To confirm that no copy of the gene was present in the genome of the transduced strain, a PCR with DreamTaq polymerase (Thermo Scientific, Dreieich, Germany) and internal primers (“int”) binding inside of the gene coding sequence was performed. The selective marker was removed by growing a culture to OD_600_ ∼ 0.2 and subsequent addition of 50 mM L-Rhamnose, followed by cultivation for ∼4h at 30 °C for flippase expression induction. Successful removal of the KmR gene from the respective locus in colonies that only grew on LB in absence of the respective antibiotic was confirmed by PCR using the locus specific KO-Ver primers and DreamTaq polymerase (Thermo Scientific, Dreieich, Germany).

#### 2.1.4. Gene overexpression in *E. coli*

Genes encoding for aspartate-glyoxylate aminotransferase (BhcA, Uniprot A1B8Z3), β-hydroxyaspartate aldolase (BhcC, Uniprot A1B8Z1), β-hydroxyaspartate dehydratase (BhcB, Uniprot A1B8Z2) and iminosuccinate reductase (BhcD, Uniprot A1B8Z0) from *Paracoccus denitrificans* were synthesized by Twist Bioscience (San Francisco, CA, USA) after codon adaptation to the codon usage of *E. coli* (Grote et al., 2005) and removal of restriction sites relevant for cloning (Zelcbuch et al., 2013). The sequences of the codon-optimized genes are given in the Supplementary Information.

Cloning was performed in *E. coli* DH5α. All genes were cloned into the pNivC vector downstream of ribosome binding site “C” (AAGTTAAGAGGCAAGA) (Zelcbuch et al., 2013) via *Mph1103*I and *Xba*I. *bhcB* was added by cutting the vector backbone pNiv-BhcC with *Nhe*I and *Xho*I and ligation with *bhcB* from pNiv-BhcB cut with *Bcu*I and *Sal*I. *bhcD* was added in the same manner with pNiv-BhcCB as accepting vector. For transfer of the genes into the expression vector pZ-ASS (p15A origin, Streptomycin resistance, strong promoter) (Braatsch et al., 2008; Wenk et al., 2018), the restriction enzymes *EcoR*I and *Pst*I (FastDigest, Thermo Scientific) were used. The genes were cloned as an operon; between the stop codon of one gene and the start codon of the next gene, there is a scar from the restriction-based cloning (TCTAGAGCTAG), a spacer sequence (TTAATAGAAATAATTTTGTTTAACTTTA) and the aforementioned ribosomal binding site. Successful ligation of insert and vector was confirmed by PCR with the primers pZ-ASS-seq-fwd and Cap-seq-rvs (Supplementary Table 2) and DreamTaq polymerase (Thermo Scientific, Dreieich, Germany). The correct sequence of constructed vectors was confirmed by Sanger sequencing of the plasmid (LGC Genomics, Berlin, DE) using the software Geneious 8 (Biomatters, New Zealand) for *in silico* cloning and sequence analysis.

#### 2.1.5. Media and growth experiments of *E. coli*

LB medium (1% NaCl, 0.5% yeast extract, 1% tryptone) (Bertani, 1951) was used for cloning, generation of deletion strains, and strain maintenance. When appropriate, kanamycin (25 μg/mL), ampicillin (100 μg/mL), streptomycin, (100 μg/mL), or chloramphenicol (30 μg/mL) were used as selective antibiotics. Growth experiments were carried out without antibiotics in standard M9 minimal medium (50 mM Na_2_HPO_4_, 20 mM KH_2_PO_4_, 1 mM NaCl, 20 mM NH_4_Cl, 2 mM MgSO_4_ and 100 μM CaCl_2_, 134 μM EDTA, 13 μM FeCl_3_·6H_2_O, 6.2 μM ZnCl_2_, 0.76 μM CuCl_2_·2H_2_O, 0.42 μM CoCl_2_·2H_2_O, 1.62 μM H_3_BO_3_, 0.081 μM MnCl_2_·4H_2_O). Carbon sources were used as indicated in the text, with the following concentrations except when stated otherwise: 20 mM glycerol, 80 mM glycolate, 10 mM glycine, 5 mM 2-oxoglutarate. For growth experiments, overnight cultures of the oxaloacetate auxotroph or the aspartate auxotroph were incubated in 4 mL M9 medium containing 20 mM glycerol and 5 mM aspartate, while overnight cultures of WT and Δ*gcl* were grown in 4 mL M9 medium containing 20 mM glycerol. Cultures were harvested by centrifugation (6,000 *g*, 3 min) and washed three times in M9 medium to remove residual carbon sources. Growth experiments were inoculated with the washed cells to an OD_600_ of 0.01 in 96-well microtiter plates (Nunclon Delta Surface, Thermo Scientific) at 37 °C. In these plates, each well contained 150 μL of culture and 50 μL mineral oil (Sigma-Aldrich) to avoid evaporation while allowing gas exchange. Growth of technical triplicates was monitored at 37 °C by absorbance measurements (600 nm) of each well every ∼10 minutes with intermittent orbital and linear shaking in a BioTek Epoch 2 plate reader (BioTek, Bad Friedrichshall, Germany). As had previously been established empirically for the instrument, blank measurements were subtracted and OD_600_ measurements were converted to cuvette OD_600_ values by multiplying with a factor of 4.35.

#### 2.1.6. Isolation and sequence analysis of Δ*gcl* pZ-ASS-BhcCBD mutant strains

The Δ*gcl* pZ-ASS-BhcCBD strain was inoculated to an OD600 of 0.01 in tube cultures of 4 mL M9 + 80 mM glycolate + 10 mM glycine. Cell growth was monitored during prolonged incubation at 37 °C for 7-20 days. Within 18 days one culture started to grow and reached an OD of 1.93. To generate single colonies, cells were dilution-streaked on LB. Isolates were inoculated into tube cultures of 4 mL M9 + 80 mM glycolate + 10 mM glycine, and the culture which grew to OD 1.97 (mutant Δ*gcl* 1) within a day was conserved and subsequently used for genome resequencing. To that end, both mutant Δ*gcl* 1 and parental Δ*gcl* pZ-ASS-BhcCBD were inoculated in LB + Strep100 medium. Of these overnight cultures, 2 ml with approx. 2 x 10^9^ cells were used for extraction of genomic DNA using the Macherey-Nagel NucleoSpin Microbial DNA purification kit (Macherey-Nagel, Düren, Germany). PCR-free libraries (microbial short insert libraries) for single-nucleotide variant detection and 150 bp paired-end reads on an Illumina HiSeq 3000 platform were constructed and sequenced by Novogene (Cambridge, UK). The reads were mapped to the reference genome of *E. coli* K-12 MG1655 (GenBank accession U00096.3) using the software Breseq (Deatherage et al., 2014). Additionally, the reads were mapped against the plasmid sequence and searched for mutations with the Find Variations/SNPs tool (minimum coverage = 20; minimum variant frequency = 50%) in Geneious. With the algorithms supplied by the software package, we identified single-nucleotide variants (with >50% prevalence in all mapped reads) and searched for regions with coverage deviating more than 2 standard deviations from the global median coverage. For generation of more mutants in a high-throughput manner, cells from a LB-Strep100 overnight culture of the Δ*gcl* pZ-ASS-BhcCBD strain were washed three times in M9 and then plated on selective M9 plates supplemented with 40 mM glycolate, 10 mM glycine and Strep100. Within 13 days, small colonies were obtained which were restreaked on LB-Strep100 plates. From these single colony restreaks, tubes with M9 + 40 mM glycolate + 10 mM glycine + Strep100 were inoculated to an OD600 of 0.01. Of 34 tubes, growth occurred in two tubes within three (mutant Δ*gcl* 2) and ten days (mutant Δ*gcl* 3), respectively. Both mutants were propagated three times in M9 + 40 mM glycolate + 10 mM glycine + Strep100. Using cells from propagation 1 (mutant Δ*gcl* 2) and propagation 2 (mutant Δ*gcl* 3) as preculture, growth dependent on glycolate and glycine was confirmed in a plate reader-based growth assay. 2 ml of mutant culture from propagation round 3 were used for gDNA isolation using the Macherey-Nagel NucleoSpin Microbial DNA purification kit (Macherey-Nagel, Düren, Germany). Whole genome sequencing was performed via the INVIEW Resequencing service for bacterial genomes up to 10 Mb with an Illumina standard genomic library provided by Eurofins Genomics (Ebersberg, Germany). The generated reads were mapped against the reference genome of *E. coli* K-12 MG1655 (GenBank accession U00096.3) using the software Breseq as described previously. Mutations found in all mutants were compared to the ones found in the parental Δ*gcl* pZ-ASS-BhcCBD strain, and all deviations from the parental genome were listed as mutant specific mutations.

#### 2.1.7. ^13^C labeling of proteinogenic amino acids

For stationary isotope tracing of proteinogenic amino acids, the oxaloacetate auxotroph + pZ-ASS-BhcCBD and a wildtype control were cultured in 4 ml of M9 medium supplemented with 5 mM unlabeled glycolate, 5 mM ^13^C_2_-labeled glycine, 20 mM glycerol, and 5 mM 2-oxoglutarate. The *Δgcl* pZ-ASS-BhcCBD mutant Δ*gcl* 1 was cultured in 4 ml of M9 medium supplemented with either unlabeled glycolate and ^13^C_2_ glycine, or ^13^C_2_ glycolate and unlabeled glycine. Cells that reached stationary growth phase were collected by centrifugation (3 min, 18,407 g) and hydrolyzed by incubation for 24 h at 95 °C with 1 ml of 6N hydrochloric acid. HCl was removed by evaporation under an air stream at 95 °C for 24 h, followed by sample resuspension in 200 µl H_2_O and removal of insoluble compounds by centrifugation (10 min, 16,000 *g*). Supernatants were used for analysis of amino acid masses by UPLC-ESI-MS with a Waters Acquity UPLC system using a HSS T3 C_18_ reversed phase column (100 mm × 2.1 mm, 1.8 μm; Waters) as described previously (Giavalisco et al., 2011). The mobile phases were 0.1 % formic acid in H_2_O (A) and 0.1% formic acid in acetonitrile (B). The flow rate was 0.4 ml/min with a gradient of 0 to 1 min – 99% A; 1 to 5 min – linear gradient from 99% A to 82%; 5 to 6 min – linear gradient from 82% A to 1% A; 6 to 8 min – kept at 1% A; 8-8.5 min – linear gradient to 99% A; 8.5-11 min – re-equilibrate. Mass spectra recorded during the first 5 min of the LC gradients were determined with an Exactive mass spectrometer (Thermo Scientific) in positive ionization mode with a scan range of 50.0 to 300.0 m/z. Data were analyzed using Xcalibur (Thermo Scientific). Amino acid standards (Sigma-Aldrich, Germany) were analyzed for determination of the retention times under the same conditions.

### 2.2. Methods for *P. putida* work (carried out at Max Planck Institute Marburg)

#### 2.2.1. Chemicals & Reagents

Unless otherwise stated, all chemicals and reagents were acquired from Merck/Sigma-Aldrich (Taufkirchen, Germany), and were of the highest purity available.

#### 2.2.2. Strains, media and cultivation conditions

All strains used in this study are listed in Supplementary Table 1. *Escherichia coli* DH5α (for genetic work) and BL21 AI (for protein production) were grown at 37 °C in lysogeny broth (Bertani, 1951). *Pseudomonas putida* KT2440 (Bagdasarian et al., 1981) and its derivatives were grown at 30 °C in lysogeny broth or in mineral salt medium (Hartmans et al., 1989) supplemented with ethylene glycol. *P. putida* transformants harboring pBG35-BhcABCD were selected on LB agar plates with 25 µg mL^-1^ tetracycline. To monitor growth, the OD_600_ of culture samples was determined on a photospectrometer (Merck Chemicals GmbH, Darmstadt, Germany).

#### 2.2.3. Vector construction

The genes encoding for the four enzymes of the BHAC from *P. denitrificans* (*bhcABCD*) were amplified in two separate fragments (*bhcAB*, *bhcCD*) using the primers given in Supplementary Table 2. The resulting PCR products were used to perform Gibson assembly (Gibson et al., 2009) with the vector backbone pTE104 that had been digested with the restriction enzyme SpeI (New England Biolabs, Frankfurt am Main, Germany) to generate pTE104-BhcABCD. The *P*coxB promoter was subsequently excised from the plasmid using XbaI and HindIII and replaced with the synthetic *P*BG35 promoter that had been generated via primer hybridization to generate pBG35-BhcABCD. Successful cloning was verified by DNA sequencing (Eurofins Genomics, Ebersberg, Germany). All plasmids used in this study are listed in Supplementary Table 1.

#### 2.2.4. Enzyme activity assays in *P. putida* cell extracts

*P. putida* cultures were harvested during mid-exponential phase (OD_600_ 0.5 – 0.7), resuspended in ice-cold 100 mM potassium phosphate buffer (pH 7.2) and lysed by sonication. Cell debris was separated by centrifugation at 35,000 x g and 4 °C for 1 h. Total protein concentration of the resulting cell-free extracts was determined by Bradford assay (Bradford, 1976) using bovine serum albumin as standard. The assays for activity of BhcA/B/C/D were performed as described previously using the relevant purified coupling enzymes (Schada von Borzyskowski et al., 2019). In all enzyme assays, the oxidation of NADH was followed at 340 nm and 30 °C on a Cary 60 UV-Vis photospectrometer (Agilent, Santa Clara, USA) in quartz cuvettes with a pathlength of 10 mm (Hellma Optik GmbH, Jena, Germany).

#### 2.2.5. High-throughput growth assays

Cultures of *P. putida* KT2440 derivatives were pre-grown at 30 °C in LB medium containing 25 µg mL^-1^ tetracycline, when appropriate. Cells were harvested, washed once with minimal medium containing no carbon source and used to inoculate growth cultures of 180 µL minimal medium containing 20 mM ethylene glycol. Growth in 96-well plates (Thermo Fisher Scientific, Waltham, USA) was monitored at 30 °C at 600 nm in a Tecan Infinite M200Pro reader (Tecan, Männedorf, Switzerland). The resulting data was evaluated using GraphPad Prism 8.1.1.

#### 2.2.6. Adaptive laboratory evolution

Three replicate populations of Δ*gcl* + BHAC were established in shake flasks and subjected to serial transfer in minimal medium containing 60 mM ethylene glycol. The cultures were inoculated to an OD_600_ of 0.01 and allowed to grow for 6-7 generations. Subsequently, the cultures were diluted 1:100 into fresh medium. The cultures were subjected to 29 transfers in total, resulting in growth for 180-200 generations. At the end of the experiment, diluted culture samples were plated on minimal medium plates containing 60 mM ethylene glycol and single colonies were isolated. These evolved isolates were cryo-conserved and used for subsequent experiments.

#### 2.2.7. Whole-cell shotgun proteomics

To acquire the proteome of parental and evolved *P. putida* strains, 20 mL cultures were grown to mid-exponential phase (OD_600_ *∼*0.4) in minimal medium supplemented with 20 mM EG. Three replicate cultures were grown for each strain. Main cultures were inoculated from precultures grown in the same medium in a 1:1,000 dilution. Cultures were harvested by centrifugation at 4,000 × g and 4 °C for 15 min. Supernatant was discarded and pellets were washed in 40 mL phosphate buffered saline (PBS; 137 mM NaCl, 2.7 mM KCl, 10 mM Na*_2_*HPO*_4_*, 1.8 mM KH*_2_*PO*_4_*, pH 7.4). After washing, cell pellets were resuspended in 1 mL PBS, transferred into Eppendorf tubes, and repeatedly centrifuged. Cell pellets in Eppendorf tubes were snap-frozen in liquid nitrogen and stored at -80 °C until they were used for the preparation of samples for LC-MS analysis and label-free quantification.

For protein extraction, bacterial cell pellets were resuspended in 2% sodium lauroyl sarcosinate (SLS) and lysed by heating (95 °C, 15 min) and sonication (Hielscher Ultrasonics GmbH, Teltow, Germany). Reduction was performed for 15 min at 90 °C in the presence of 5 mM tris(2-carboxyethyl)phosphine (TCEP) followed by alkylation using 10 mM iodoacetamide at 25 °C for 30 min. The protein concentration in each sample was determined using the BCA protein assay kit (Thermo Fisher Scientific, Waltham, USA) following the manufacturer’s instructions. Protein cleanup and tryptic digest were performed using the SP3 protocol as described previously (Moggridge et al., 2018) with minor modifications regarding protein digestion temperature and solid phase extraction of peptides. SP3 beads were obtained from GE Healthcare (Chicago, USA). 1 µg trypsin (Promega, Fitchburg, USA) was used to digest 50 μg of total solubilized protein from each sample. Tryptic digest was performed overnight at 30 °C. Subsequently, all protein digests were desalted using C18 microspin columns (Harvard Apparatus, Holliston, USA) according to the manufacturer’s instructions.

LC-MS/MS analysis of protein digests was performed on a Q-Exactive Plus mass spectrometer connected to an electrospray ion source (Thermo Fisher Scientific, Waltham, USA). Peptide separation was carried out using an Ultimate 3000 nanoLC-system (Thermo Fisher Scientific, Waltham, USA), equipped with an in-house packed C18 resin column (Magic C18 AQ 2.4 µm; Dr. Maisch, Ammerbuch-Entringen, Germany). The peptides were first loaded onto a C18 precolumn (preconcentration set-up) and then eluted in backflush mode with a gradient from 98% solvent A (0.15% formic acid) and 2% solvent B (99.85% acetonitrile, 0.15% formic acid) to 25% solvent B over 48 min, continued from 25% to 35% of solvent B for an additional 34 min. The flow rate was set to 300 nL/min. The data acquisition mode for the initial LFQ study was set to obtain one high-resolution MS scan at a resolution of 60,000 (*m/z* 200) with scanning range from 375 to 1500 *m/z* followed by MS/MS scans of the 10 most intense ions. To increase the efficiency of MS/MS shots, the charged state screening modus was adjusted to exclude unassigned and singly charged ions. The dynamic exclusion duration was set to 30 sec. The ion accumulation time was set to 50 ms (both MS and MS/MS). The automatic gain control (AGC) was set to 3 × 10^6^ for MS survey scans and 1 × 10^5^ for MS/MS scans. Label-free quantification was performed using Progenesis QI (version 2.0). MS raw files were imported into Progenesis and the output data (MS/MS spectra) were exported in mgf format. MS/MS spectra were then searched using MASCOT (version 2.5) against a database of the predicted proteome from *P. putida* KT2440 downloaded from the UniProt database (www.uniprot.org; download date 03/01/2021), containing 386 common contaminant/background proteins that were manually added. The following search parameters were used: full tryptic specificity required (cleavage after lysine or arginine residues); two missed cleavages allowed; carbamidomethylation (C) set as a fixed modification; and oxidation (M) set as a variable modification. The mass tolerance was set to 10 ppm for precursor ions and 0.02 Da for fragment ions for high energy-collision dissociation (HCD). Results from the database search were imported back to Progenesis, mapping peptide identifications to MS1 features. The peak heights of all MS1 features annotated with the same peptide sequence were summed, and protein abundance was calculated per LC–MS run. Next, the data obtained from Progenesis were evaluated using the SafeQuant R-package version 2.2.2 (Glatter et al., 2012). Hereby, 1% FDR of identification and quantification as well as intensity-based absolute quantification (iBAQ) values were calculated.

#### 2.2.8. DNA extraction, library preparation, Illumina sequencing

Genomic DNA extraction from the parental and evolved strains of *P. putida* KT2440 Δ*gcl* + BHAC was performed with the Nucleo®Spin Microbial DNA kit (Macherey Nagel, Düren, Germany). Eppendorf DNA LoBind Tubes were used at all times. DNA concentrations were measured using the Qubit dsDNA BR Assay Kit (Thermo Fischer Scientific, Waltham, USA). Libraries were prepared using 250 ng of isolated genomic DNA following the NEBNext® Ultra™ II FS DNA Library Prep (New England Biolabs, Frankfurt am Main, Germany) with Sample Purification Beads (NEB E6177S). NEBNext® Multiplex Oligos for Illumina® (NEB E7500) were used. An approximate insert size distribution of 200-350 bps was selected using the provided beads according to manufacturer’s instructions. Adaptor-ligated DNA was enriched through 4 cycles of PCR. Quality and size distribution of the libraries was assessed on a Bioanalyzer High Sensitivity DNA Analysis chip on an Agilent 2100 Bioanalyzer (Agilent, Waldbronn, Germany). Prepared libraries were sequenced with the MiniSeq High Output Reagent Kit (300-cycles) (Illumina, Berlin, Germany) on a MiniSeq system in paired-end mode (2 x 150 bp). Sequence analysis was carried out using Geneious Prime 2020.1.2 (Biomatters, Auckland, New Zealand). Reads were mapped against the *P. putida* KT2440 genome using GenBank Accession file AE015451 and pBG35-BhcABCD (BHAC expression plasmid) as reference replicons. Variants were detected with the Find Variations/SNPs tool (minimum coverage = 20; minimum variant frequency = 95%) and compared to variants detected in parental control reads mapped to the same reference replicons.

## 3. Results

### Developing the BHA shunt for prototyping the core reactions of the BHAC in *E. coli*

The core reaction sequence of the BHAC consists of three enzymes, BHA aldolase (BhcC), BHA dehydratase (BhcB), and iminosuccinate reductase (BhcD). Together, these three enzymes convert glyoxylate and glycine into aspartate, constituting a linear metabolic module that we termed BHA shunt (Figure 1d). To probe whether the BHA shunt would work in a gammaproteobacterial host, we designed three *E. coli* selection strains with different auxotrophies. These strains show increasing demand for flux through the BHA shunt, starting from a specific amino acid auxotrophy (aspartate auxotroph), over central carbon metabolism auxotrophy (oxaloacetate auxotroph) to biosynthesis of the complete biomass (glyoxylate caboligase (*gcl*) deletion strain) from glycine and glycolate. We chose glycolate over glyoxylate, the actual substrate of BhcC, as the latter is a reactive aldehyde that can be detrimental to cells in higher concentrations. *E. coli* converts glycolate into glyoxylate via glycolate oxidase (Pellicer et al., 1996).

### The BHA shunt complements an aspartate auxotrophic *E. coli* strain

We first created an aspartate auxotrophic strain as described previously (Gelfand et al., 1977) by deleting the genes *aspC*, encoding for aspartate aminotransferase, and *tyrB*, encoding for aromatic amino acid aminotransferase (Figure 2a). Both of these enzymes catalyze the transfer of an amino group from glutamate to oxaloacetate for aspartate biosynthesis (Hayashi et al., 1993; Reitzer, 2014), making the double mutant auxotrophic for aspartate and amino acids of the aspartate family (asparagine, lysine, methionine, threonine, isoleucine). Note that the aspartate auxotroph is only incapable of oxaloacetate amination, while the TCA cycle and other central metabolic pathways are still fully functional in this strain.

**Figure 2:**
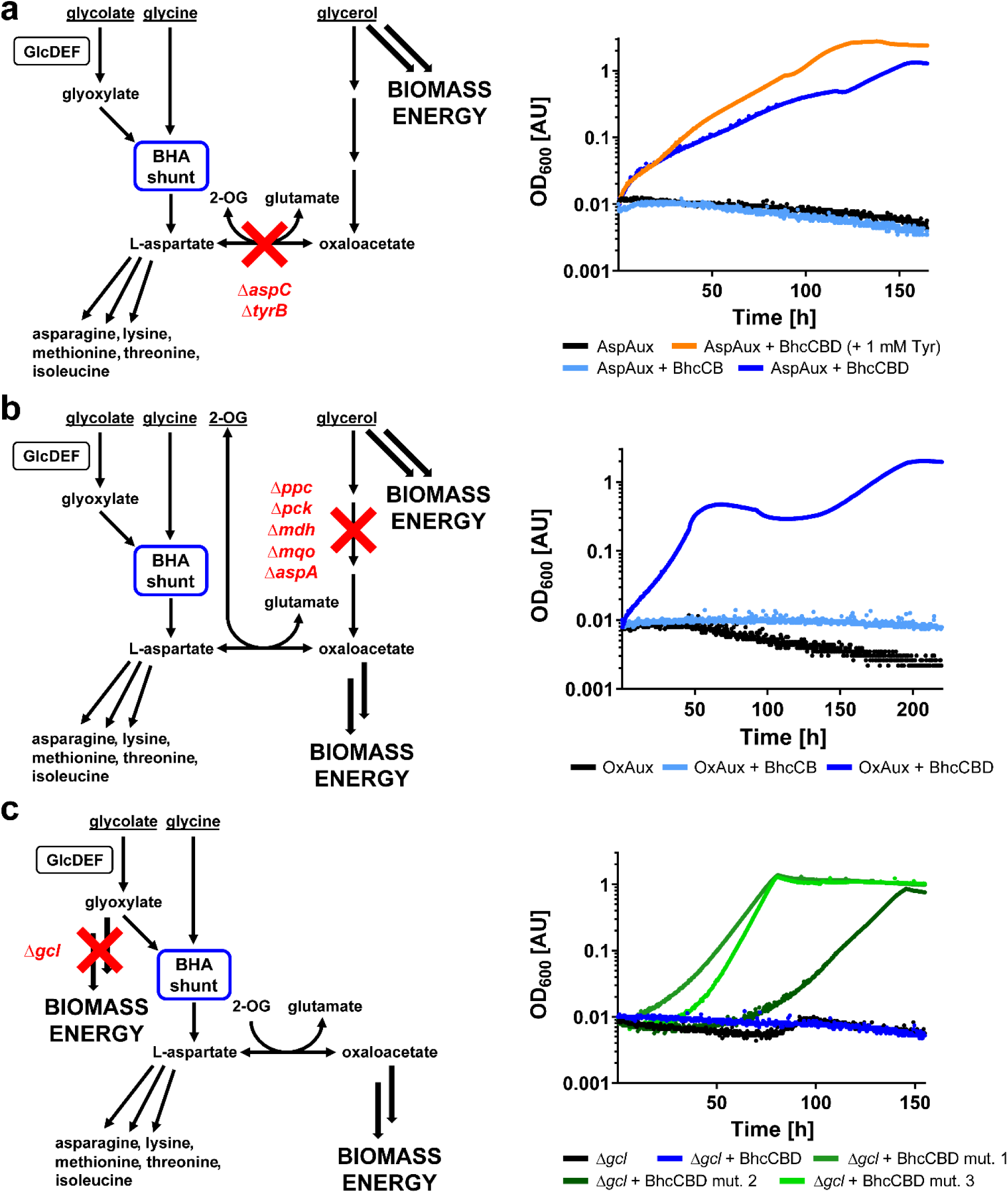
Selection schemes and growth assays of *E. coli* auxotrophic strains engineered with the BHA shunt. Compounds that are underlined in the schemes were provided in the growth medium. **a**, The aspartate auxotrophic strain cannot aminate oxaloacetate to generate aspartate due to deletion of *aspC* and *tyrB*. When expressing the BHA shunt, this strain is able to grow in the presence of glycerol, glycolate, and glycine (blue growth curve). Supplementation of 1 mM tyrosine to the medium results in improved growth (orange). In contrast, the same strain without the BHA shunt (black) or with a plasmid expressing only *bhcC* and *bhcB* (light blue) is unable to grow. **b**, The oxaloacetate auxotrophic strain lacks all enzymes producing oxaloacetate via the TCA cycle or anaplerosis due to deletion of *ppc*, *pck*, *mdh*, *mqo* and *aspA*. When expressing the BHA shunt, this strain is able to grow in the presence of glycerol, glycolate, glycine, and 2-oxoglutarate (blue). In contrast, the same strain without the BHA shunt (black) or with a plasmid expressing only *bhcC* and *bhcB* (light blue) is unable to grow. **c**, The Δ*gcl* strain cannot utilize glyoxylate via the glycerate pathway. When expressing the BHA shunt, this strain was not able to grow in the presence of glycolate and glycine (blue), similar to the same strain without the BHA shunt (black). However, three isolated mutants of Δ*gcl* expressing the BHA shunt were able to grow on those carbon sources (shades of green). In all cases, representative growth curves from n ≥ 3 independent cultures are shown, with errors < 5%.

When growing with glycerol as carbon source together with glycolate and glycine, the mutant expressing the BHA shunt immediately grew with a doubling time of 18.2 ± 0.4 h, while negative control strains expressing an incomplete BHA shunt (without *bhcD*) or carrying an empty plasmid did not grow (Figure 2a). The slow growth of the strain is likely due to tyrosine limitation caused by the *tyrB* knockout, which can be only partially complemented by the branched-chain amino acid aminotransferase IlvE (Lee-Peng et al., 1979). Indeed, addition of 1 mM tyrosine to the medium improved growth of the aspartate auxotroph with the BHA shunt (SI Figure 1), resulting in a doubling time of 13 ± 0.4 h. Altogether, these results showed that the BHA shunt was generally active in *E. coli*.

### The BHA shunt complements an oxaloacetate auxotrophic *E. coli* strain

Next, we created a strain that is auxotrophic for oxaloacetate, which is required both for the biosynthesis of aspartate and amino acids of the aspartate family, as well as for fueling the TCA cycle. We deleted the genes encoding for phosphoenolpyruvate carboxylase (Ppc), phosphoenolpyruvate carboxykinase (Pck), malate dehydrogenase (Mdh), malate:quinone oxidoreductase (Mqo), and aspartate ammonia-lyase (AspA) to suppress oxaloacetate synthesis in *E. coli* (Figure 2b). This strain would only be able to grow if both aspartate and oxaloacetate are formed through the BHA shunt, the latter either through spontaneous hydrolysis of iminosuccinate or through amino transfer from aspartate to 2-oxoglutarate (2-OG) by the previously mentioned AspC and TyrB.

However, we did not observe growth of the oxaloacetate auxotroph on glycerol, glycine, and glycolate or glyoxylate (SI Figure 2), unless we provided additionally 5 mM 2-OG (Figure 2b). Furthermore, only the oxaloacetate auxotroph expressing the full BHA shunt grew on glycerol, glycine, glycolate and 2-OG, while an oxaloacetate auxotroph lacking iminosuccinate reductase couldn’t grow in the same conditions (Fig. 2b). When tested on glycerol and 2-OG alone, the oxaloacetate autotroph did not grow (SI Figure 2), indicating that growth of the oxaloacetate auxotroph indeed required both, a functional BHA shunt and 2-OG. We concluded that 2-OG addition was necessary to provide the amino group acceptor for glutamate and glutamate family amino acids, and to facilitate oxaloacetate synthesis. Since expression of iminosuccinate reductase was required for growth of the oxaloacetate auxotroph, we suggest that no relevant spontaneous hydrolysis of iminosuccinate into oxaloacetate took place *in vivo* (Mortarino et al., 1996; Schada von Borzyskowski et al., 2019). We used ^13^C-labeling to follow the fate of the different carbon sources *in vivo*. When growing the oxaloacetate auxotrophic strain with the BHA shunt on glycerol, 2-OG, glycolate, and fully ^13^C-labeled glycine, aspartate and threonine were double-labeled, in line with the hypothesis that half of the backbone was derived from ^13^C-labeled glycine, while the other half was derived from unlabeled glycolate. Methionine was triple-labeled in line with the hypothesis that the methyl-group was derived from ^13^C-labeled glycine via the glycine cleavage complex. These results (Figure 3a) demonstrated that the three amino acids were indeed synthesized via the BHA shunt. In contrast, proline was unlabeled, demonstrating that it was solely synthesized from 2-OG via glutamate. In summary, the ^13^C-labeling underlined that the BHA shunt is active in the oxaloacetate auxotroph, but that addition of 2-OG is required as precursor for the production of glutamate family amino acids.

**Figure 3:**
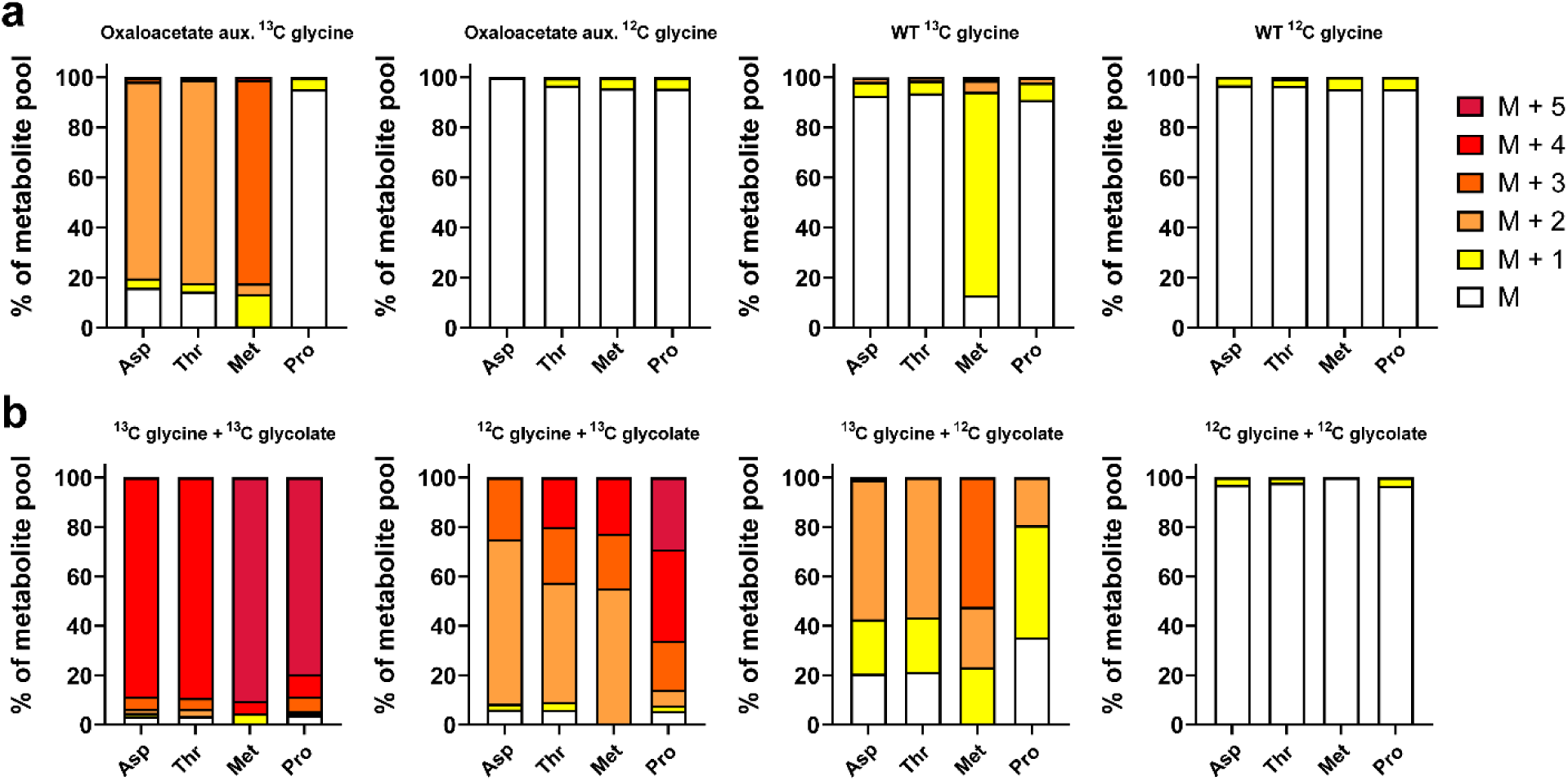
^13^C-labeling of selected amino acids in auxotrophic *E. coli* strains expressing the BHA shunt. **a**, ^13^C-labeling of aspartate, threonine, methionine, and proline in the oxaloacetate auxotroph strain with the BHA shunt or a WT control grown on glycerol, 2-OG, glycolate, and either fully ^13^C-labeled or unlabeled glycine. **b**, ^13^C-labeling of aspartate, threonine, methionine, and proline in mutant Δ*gcl* 1 with the BHA shunt grown on different combinations of fully ^13^C-labeled and unlabeled glycolate and glycine.

### The BHA shunt provides all biomass in an evolved *E. coli* Δ*gcl* strain

Finally, we sought to test whether the BHA shunt would be able to sustain complete biomass formation in *E. coli*. To this end, we used a selection strain, in which *gcl*, the gene encoding for glyoxylate carboligase, the key enzyme of the glycerate pathway, is deleted. This strain cannot grow on glycolate or glyoxylate anymore, unless the BHA shunt is able to complement the *gcl* deletion (Figure 2c).

When inoculating the Δ*gcl* strain + BHA shunt in M9 medium with glycolate and glycine, we did not observe any growth. However, after prolonged incubation, we were able to isolate a spontaneous mutant (mutant Δ*gcl* 1) that was capable of growth on the two carbon sources (Figure 2c). Mutant Δ*gcl* 1 was not able to grow on M9 with glycine only, excluding the possibility that the combined activities of the glycine cleavage system and serine-hydroxymethyl transferase would permit growth. The evolved strain also did not grow on glycolate as sole carbon source, confirming that no latent glyoxylate assimilation pathway is present in *E. coli* (SI Figure 3). The ^13^C-labelling pattern using different combinations of labeled glycolate and glycine independently confirmed that the BHA shunt was responsible for growth of this mutant (Figure 3b).

We isolated two more mutants (mutants Δ*gcl* 2 and 3), which were also capable to grow with the BHA shunt (Fig. 2c). Whole genome sequencing showed that all evolved strains showed mutations in the TCA cycle. In evolved strains Δ*gcl* 2 and Δ*gcl* 3, *sucC*, the gene encoding for the β-subunit of succinyl-CoA synthetase was mutated (an insertion resulting in a frameshift in mutant Δ*gcl* 2, and a nonsense mutation in mutant Δ*gcl* 3), while Δ*gcl* 1 showed a mutation in succinate dehydrogenase. Furthermore, Δ*gcl* 1 showed three additional mutations, which seemed not directly related to central carbon metabolism.

Altogether, these experiments showed that in principle the whole biomass of *E. coli* can be generated through enzymes of the BHA shunt and that the native metabolic network of *E. coli* required only slight adaptations. Encouraged by these results, we next sought to exploit the complete four-enzyme BHAC for assimilation of EG in *P. putida*.

### Implementation of the BHAC in *P. putida* enables improved EG assimilation

Having demonstrated that it is possible to generate all biomass through the BHA shunt in *E. coli*, we next aimed at implementing the full BHAC into *P. putida*. We used a broad host range expression vector (Schada von Borzyskowski et al., 2015) to express the *bhc* gene cluster and exchanged its original promoter with the defined synthetic promoter *P*BG35 (Zobel et al., 2015). When introduced into *P. putida* KT2440 Δ*gcl*, which is not able to grow with EG as sole carbon source, the strain immediately started to grow. Quantification of enzyme activities in cell-free extracts confirmed that the heterologous BHAC enzymes were functional, at activities between 0.5 and 10 U/mg (Figure 4a).

**Figure 4:**
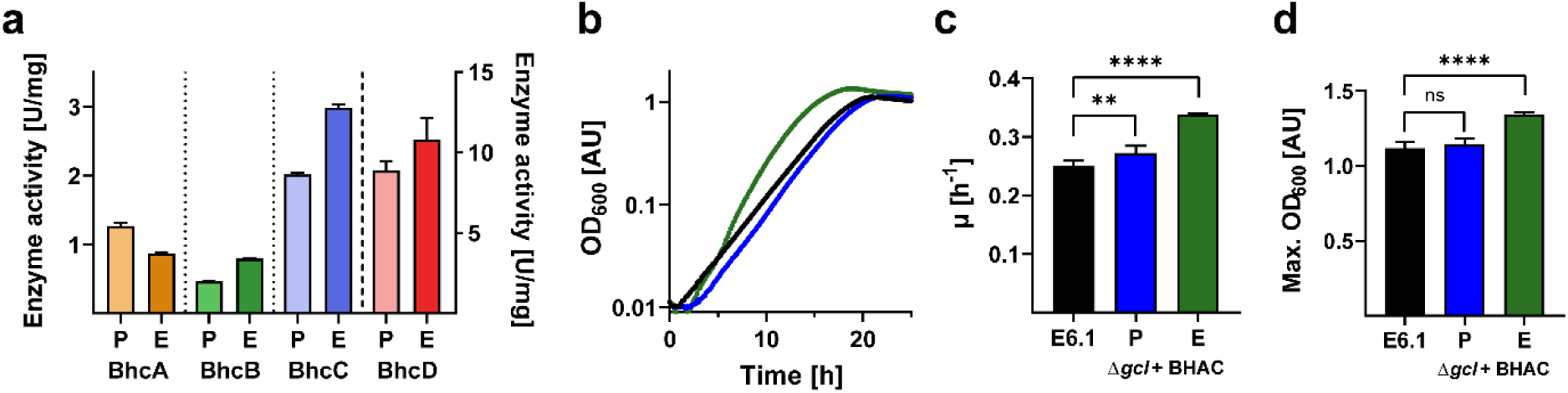
Characterization of *P. putida* KT2440 Δ*gcl* + BHAC and an evolved derivative. **a**, Specific activities of BHAC enzymes in cell-free extracts of EG-grown Δ*gcl* + BHAC. The bars labeled ‘P’ correspond to the parental engineered strain, while the darker bars labeled ‘E’ correspond to an evolved strain isolated after ALE on EG. Note that the activity of BhcD is plotted on the right y axis. Data are the mean ± s.d. of n = 3 replicate experiments. **b**, Growth curves of E6.1 (black), Δ*gcl* + BHAC (blue), and its evolved derivative (green) grown in the presence of 20 mM EG. Shown are representative growth curves from n = 6 independent cultures, with errors < 5%. **c**, Growth rates (µ) and **d**, Max. OD_600_ of E6.1 and Δ*gcl* + BHAC (parental and evolved) grown in the presence of 20 mM EG. Data are the mean ± s.d. of n = 6 independent cultures, and results were compared using an unpaired t test with Welch’s correction in GraphPad Prism 8.1.1 (****, p < 0.0001; **, p < 0.01; ns, not significant).

When growing on EG, *P. putida* KT2440 Δ*gcl* + BHAC notably behaved similar to *P. putida* KT2440 E6.1, a strain that was recently evolved to grow on EG with the glycerate pathway (Li et al., 2019) (Figure 4b). While the yield, measured as maximum OD_600_, was comparable between the two strains (Figure 4d), *P. putida* KT2440 Δ*gcl* + BHAC grew at 10% increased growth rate (Figure 4c). In summary, these results showed that the strain expressing the BHAC instantly compared to the evolved E6.1 strain with the glycerate pathway, without any optimization of promoter, ribosome binding sites, or gene sequences.

### Proteome analysis reveals changes in central carbon metabolism of *P. putida* upon introduction of the BHAC

We next were interested in understanding how the heterologously expressed BHAC affected the genetic and metabolic network in *P. putida* KT2440 Δ*gcl* + BHAC compared to strain E6.1 (Figure 5). To that end, we analyzed the proteome of both strains grown on EG.

**Figure 5:**
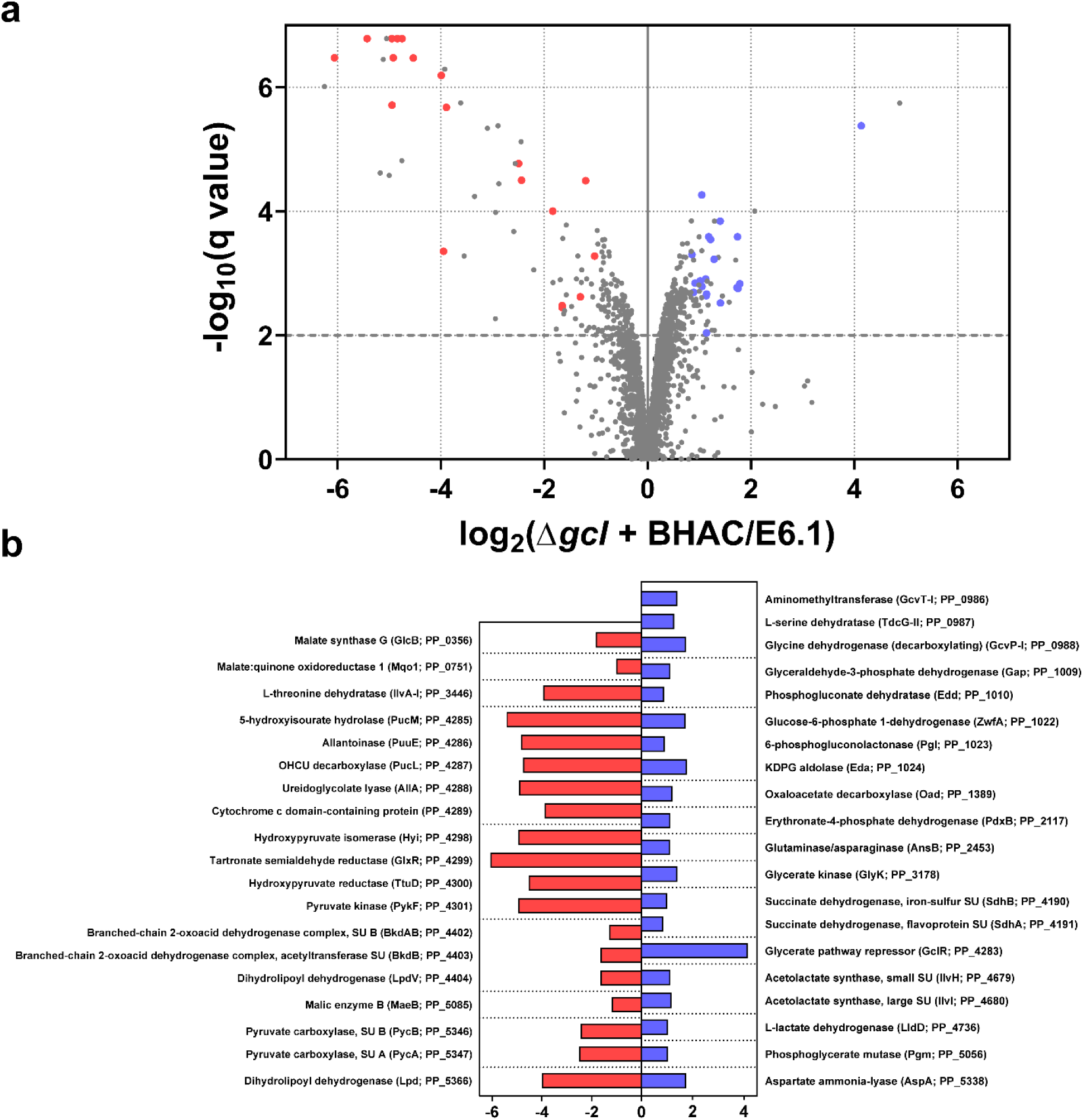
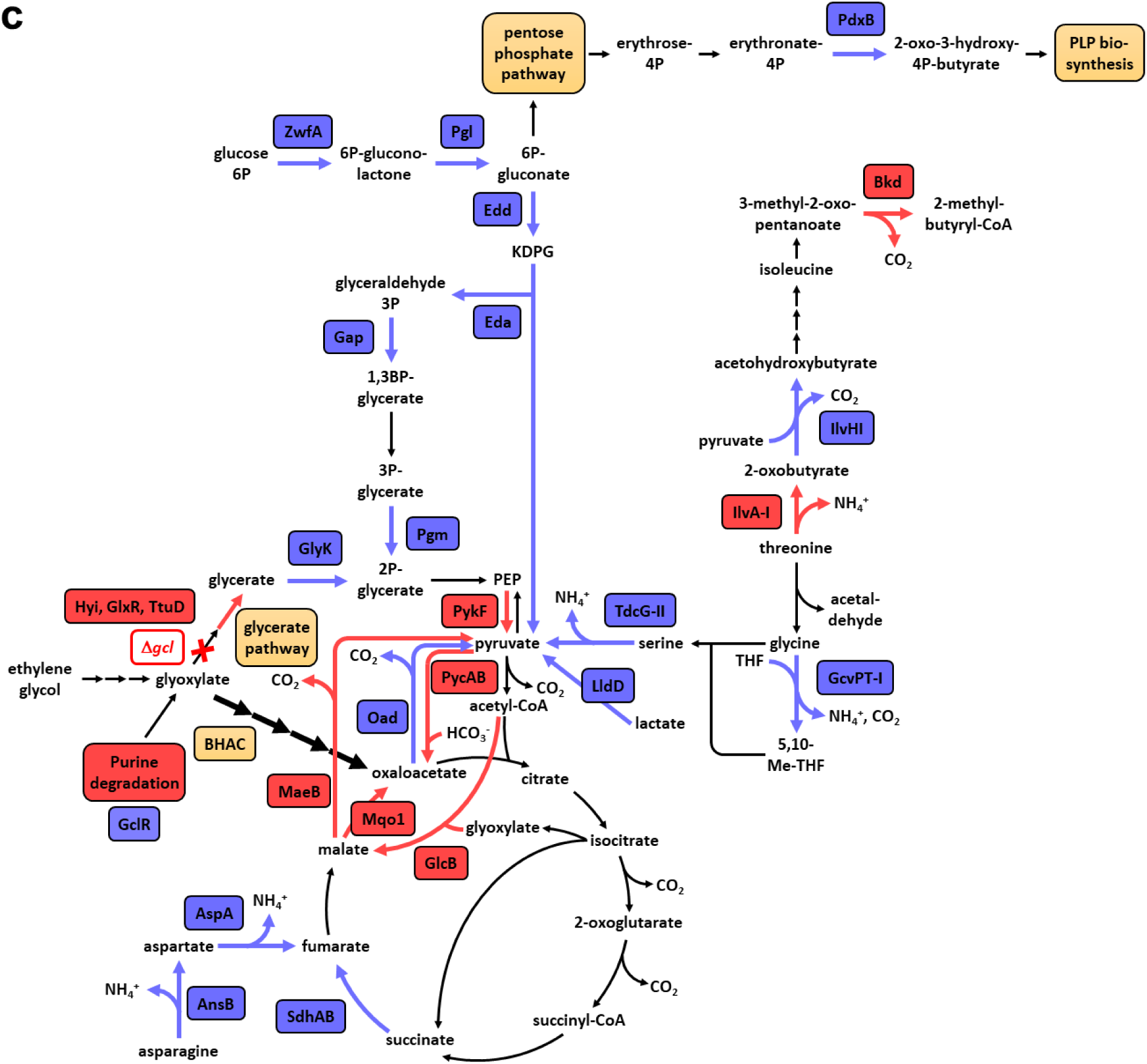
Proteome analysis of *P. putida* KT2440 Δ*gcl* + BHAC. **a**, Analysis of the proteome of EG-grown Δ*gcl* + BHAC compared to E6.1. All proteins quantified by at least three unique peptides are shown, and the proteins involved in central carbon metabolism that showed the strongest decrease or increase in abundance are marked in red or blue in the volcano plot, respectively. **b**, The log2 fold change of these proteins, sorted by locus name (in brackets). **c**, The role of these up- and downregulated proteins in central carbon metabolism of *P. putida* KT2440. Altered enzyme production levels in key metabolic routes, such as the TCA cycle, the C3-C4 node, and several amino acid biosynthesis pathways demonstrate marked changes upon introduction of the heterologous BHAC.

Significant differences between the two strains were observed on the level of enzymes in central carbon metabolism, in particular the C3-C4 node that connects and balances glycolysis and gluconeogenesis. Pyruvate carboxylase (PycAB), malate:quinone oxidoreductase (Mqo1), and malic enzyme B (MaeB) levels were all decreased in Δ*gcl* + BHAC compared to E6.1. In contrast, oxaloacetate decarboxylase (Oad) levels were increased. Decreased PycAB levels in the BHAC strain are in line with the notion that EG assimilation via the glycerate pathway produces 2-phosphoglycerate (2-PG), a glycolysis intermediate, which requires PycAB to anaplerotically feed the TCA cycle, while the BHAC yields oxaloacetate, thus directly feeding the TCA cycle (without the need for PycAB). Similarly, Mqo1, which catalyzes the quinone-dependent oxidation of malate to oxaloacetate (Koendjbiharie et al., 2021), might become obsolete in the strain carrying the BHAC, as the concentration of oxaloacetate is already much higher in the BHAC strain compared to the E6.1 strain. The observed increased levels of Oad in the BHAC strain support this notion and suggest that the enzyme acts as a valve to funnel excess oxaloacetate to pyruvate (Klaffl et al., 2010).

On the level of the TCA cycle and glyoxylate shunt, we observed an increase in succinate dehydrogenase (SdhAB) levels in the BHAC strain, while malate synthase (GlcB) was decreased. The latter is probably explained by direct feeding of the BHAC into the TCA cycle, which makes malate supply via the glyoxylate shunt obsolete in the BHAC strain.

Furthermore, enzyme levels in aspartate metabolism (AspA, AnsB) were increased in the BHAC strain, suggesting that part of the aspartate pool is deaminated to feed fumarate into the TCA cycle. Increased enzyme levels in glycine metabolism (GcvPT-I, TdG-II) indicated that some of the glyoxylate-derived glycine is further used to generate serine and pyruvate via the glycine cleavage complex and serine dehydratase in the BHAC strain. Interestingly, the levels of erythronate 4-phosphate dehydrogenase (PdxB) were also increased. Since PdxB is involved in the biosynthesis of pyridoxal 5-phosphate (PLP), the cofactor of three enzymes of the BHAC (BhcA, BhcB, BhcC), it is tempting to speculate that its upregulation provides sufficient PLP supply for growth.

As expected, glycerate pathway enzyme levels (Hyi, GlxR, TtuD) as well as pyruvate kinase (PykF), which is part of the same operon (Franden et al., 2018), were strongly decreased in Δ*gcl* + BHAC compared to E6.1. It was previously reported that E6.1 was able to grow on EG due to a mutation that resulted in truncation of the regulator GclR (Li et al., 2019). This was verified by our proteomics data, which additionally showed that (truncated) GclR levels were decreased in the E6.1 strain compared to GclR in the Δ*gcl* + BHAC strain. Furthermore, enzyme levels of a purine/allantoin degradation pathway (PP_4285-89) were also decreased, which can be explained by the fact that the corresponding genes are part of the GclR regulon (Li et al., 2019).

Finally, we observed increased levels of enzymes of the Entner-Doudoroff pathway (Edd, Eda) and several other glycolytic enzymes (ZwfA, Pgl, Gap, Pgm) in Δ*gcl* + BHAC. However, these changes are not easy to rationalize, as glycolytic pathways in *P. putida* KT2440 form a complex network (Nikel et al., 2015), and there are three different isoforms of Zwf with different cofactor specificities (Volke et al., 2021).

In summary, distinct changes of the native metabolic network of *P. putida* KT2440 resulted in immediate growth of Δ*gcl* + BHAC on EG. However, we speculated that the true potential of the BHAC was still masked and that more adaptations of the host would be required for a better integration of the heterologous module into the metabolic network of the cell.

### Evolution and characterization of a *P. putida* + BHAC strain with improved growth performance on EG

Next, we aimed at further improving growth of *P. putida* KT2440 Δ*gcl* + BHAC via adaptive laboratory evolution (ALE). We established three replicate populations in shake flasks and subjected them to serial transfer in minimal medium containing 60 mM EG. After about 200 generations, we isolated individual evolved strains and compared their growth on EG to their parental strain (SI Figure 4). One evolved isolate that exhibited the fastest growth and highest final OD was selected for more in-depth analysis. Notably, the evolved Δ*gcl* + BHAC strain showed an improved growth rate of 25% and 35% to the ancestral and the E6.1 strain, respectively, while the biomass yield, measured as final OD_600_, was improved by 18% and 20%, respectively (Figure 4). Plasmid sequencing confirmed that the *bhcABCD* expression module was unchanged in the evolved strain. Quantification of BHAC enzyme activities in cell-free extracts showed no drastic differences compared to the parental strain (Figure 4a). While activities of BhcB, BhcC, and BhcD were slightly increased in the evolved strain, BhcA activity was slightly decreased. However, these observed changes were of rather subtle nature, suggesting that the major changes allowing for improved growth of the evolved strain occurred on the level of the native metabolic and genetic networks of *P. putida* KT2440 Δ*gcl*.

Whole genome sequencing of the evolved Δ*gcl* + BHAC strain confirmed deletion of *gcl* and revealed five mutations, compared to the parental strain (Table 2). We observed a point mutation in a LysR family transcriptional regulator of unknown function (PP_1861), as well as a point mutation and a 227 bp deletion in *flhB*, encoding for a substrate specificity protein of the flagellin export apparatus. Furthermore, we also noted a point mutation in LldR, the transcriptional regulator of the lactate utilization gene cluster (Gao et al., 2012), as well as 6 bp deletion in *regB*, encoding for a sensor histidine kinase (Fernandez-Pinar et al., 2008).

**Table 1:** Mutations found in genome resequencing data of three isolated mutants of E. coli Δgcl with the BHA shunt.

| Strain | Gene | Mutation | Annotation | Position | Protein |
| --- | --- | --- | --- | --- | --- |
| $\Delta gcl$ 1 | *sdhA* | G → T | G179C | 755,665 | succinate dehydrogenase, flavoprotein subunit |
| | *kgtP* | A → T | L238I | 2,724,682 | 2-OG:H<sup>+</sup> symporter |
| | *yhiM* | (A)<sub>6</sub> → 5 | coding (1050/1053 nt) | 3,635,538 | inner membrane protein with role in acid resistance |
| | *entF* | G → A | E499E | 614,877 | apo-serine activating enzyme |
| $\Delta gcl$ 2 | *sucC* | +G | coding (1161/1167 nt) | 763,398 | succinyl-CoA synthetase, β-subunit |
| $\Delta gcl$ 3 | *sucC* | C → T | Q380* | 763,375 | succinyl-CoA synthetase, β-subunit |

**Table 2:** Mutations found in genome resequencing data of an isolated mutant of P. putida KT2440 Δgcl + BHAC.

| Gene ID | Mutation | Annotation | Position | Protein |
| --- | --- | --- | --- | --- |
| PP_0887 | ΔCCTGCG | ΔR100L101 | 1,030,236 | Sensor histidine kinase RegB |
| PP_1861 | C → T | G65S | 2,083,041 | Transcriptional regulator, LysR family |
| PP_4352 | T → A | I175F | 4,946,058 | flagellin export apparatus, substrate specificity protein FlhB |
| PP_4352 | Δ227 bp | | 4,946,139 | flagellin export apparatus, substrate specificity protein FlhB |
| PP_4734 | T → C | E242G | 5,382,556 | DNA-binding transcriptional dual regulator LldR |

LldR controls expression of the genes *lldD* (encoding for L-lactate dehydrogenase) and *dld2* (encoding for D-lactate dehydrogenase) in *P. aeruginosa* (Gao et al., 2012). Proteomics data showed that in the evolved strain, the levels of both lactate dehydrogenases were increased (Figure 6), indicating that the balance between lactate and pyruvate, and thus cellular redox management, is fine-tuned by the *lldR* mutation in the evolved Δ*gcl* + BHAC strain.

**Figure 6:**
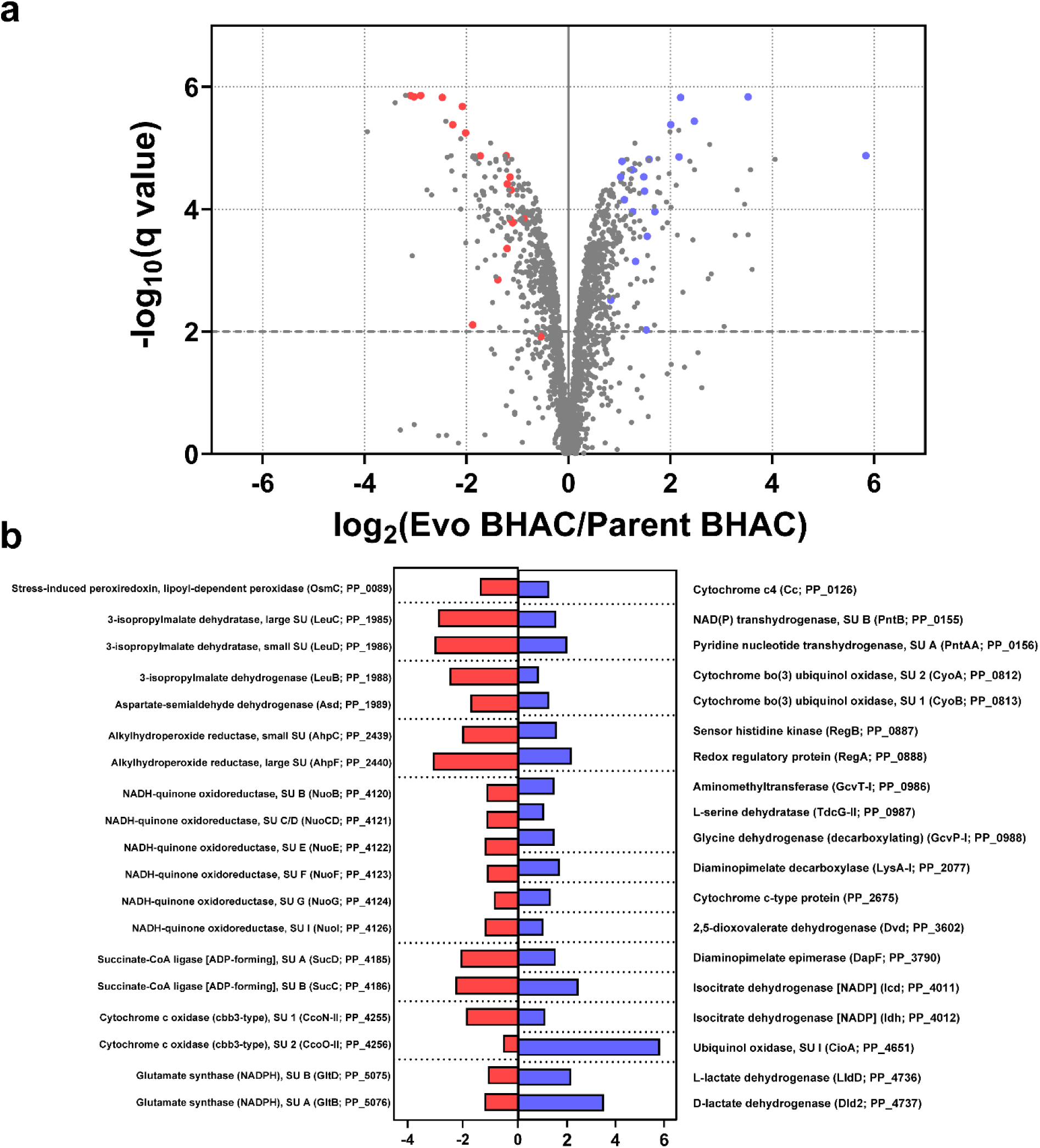
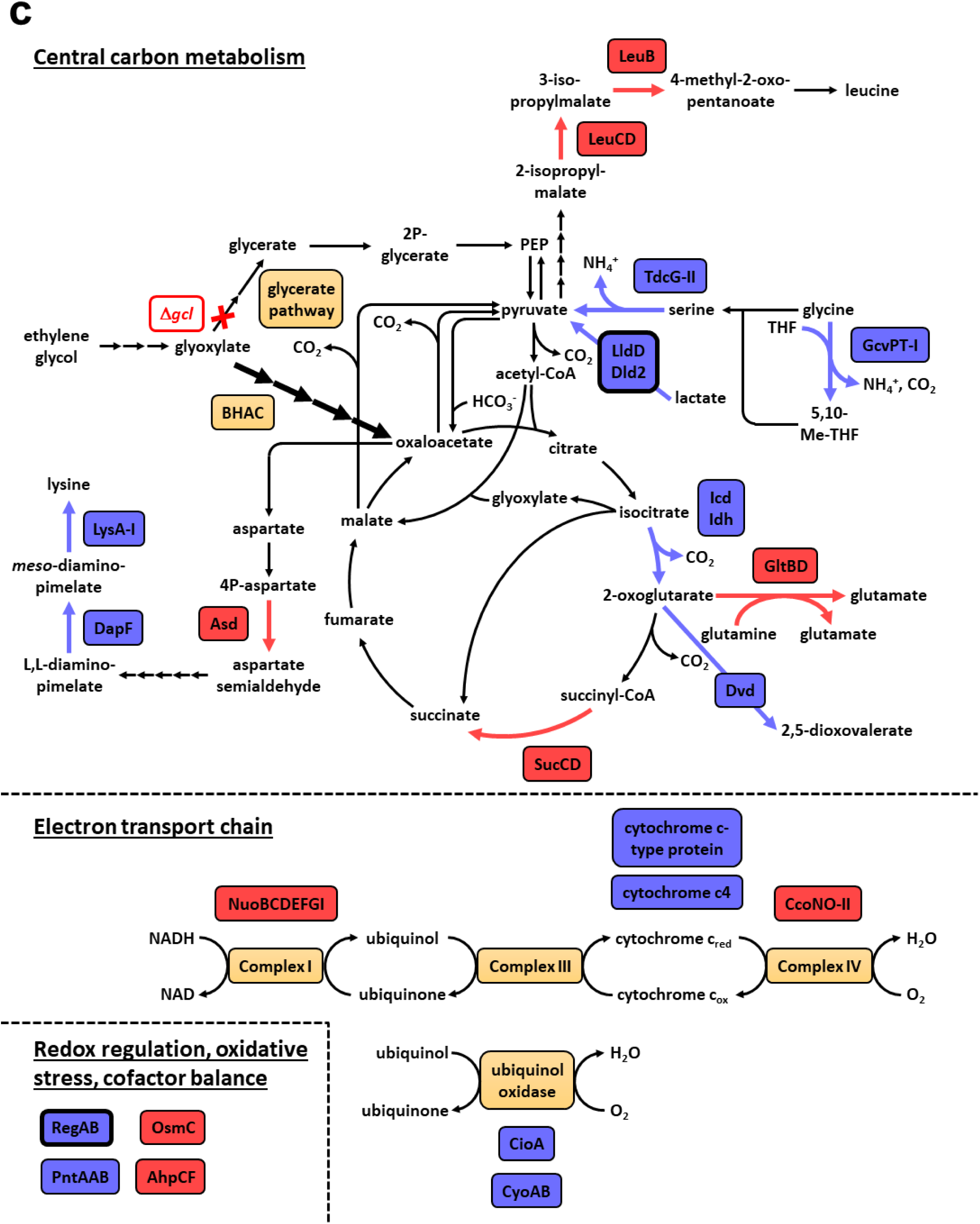
Proteome analysis of *P. putida* KT2440 Δ*gcl* + BHAC after ALE on EG. **a**, Analysis of the proteome of evolved Δ*gcl* + BHAC compared to its parental strain. All proteins quantified by at least three unique peptides are shown, and the enzymes and regulators involved in central carbon and energy metabolism that showed the strongest decrease or increase in abundance are marked in red or blue in the volcano plot, respectively. **b**, The log2 fold change of these proteins, sorted by locus name (in brackets). **c**, The role of these up- and downregulated proteins in central carbon and energy metabolism of *P. putida* KT2440. Enzyme production levels in the TCA cycle and several amino acid biosynthesis pathways, as well as in the electron transport chain, demonstrate marked changes upon prolonged growth on EG via the heterologous BHAC. Changes in protein production that are connected to mutations in the evolved strain are denoted by a black frame (LldD + Dld2, RegAB).

Redox management was likely also changed by the 6-bp *regB* deletion. RegAB is part of a conserved two-component regulatory system that regulates redox control in a wide variety of photosynthetic and non-photosynthetic bacteria (Elsen et al., 2004) (Fernandez-Pinar et al., 2008). Proteomics showed that RegA and RegB levels were increased in the evolved Δ*gcl* + BHAC strain. In *P. aeruginosa*, the homologous regulatory system RoxRS interacts with the promoter region of *cioAB*, the genes encoding for cyanide-insensitive ubiquinol oxidase, regulating their expression (Comolli & Donohue 2002). Interestingly, CioA was the most upregulated protein in the evolved Δ*gcl* + BHAC strain, suggesting that the *regB* mutation directly affected the composition of the ETC. Proteomics identified additional changes in the ETC proteome. The levels of six subunits of NADH ubiquinone oxidoreductase (complex I), as well as two subunits of a a cytochrome c oxidase were decreased, while CyoAB, another ubiquinol oxidase, and a cytochrome c-type protein (PP_2675) as well as cytochrome c4 were increased. Even though we note that subunits of ETC complexes are membrane-bound proteins, which are more difficult to quantify, our data nevertheless suggests that the ETC was partly remodeled in the evolved Δ*gcl* + BHAC strain. This remodeling is likely caused by an increased influx of ubiquinol, derived from PedE/H-mediated oxidation of EG. Our data also suggests that the majority of electrons are directly transferred to oxygen as terminal electron acceptor by the two ubiquinol oxidases, while a smaller share is shuttled to cytochromes, which are subsequently re-oxidized. Concomitant with this hypothesis, the levels of stress-induced peroxiredoxin OsmC and alkyl hydroperoxide reductase AhpCF were decreased.

Finally, the proteomics data showed further changes in central carbon metabolism in the evolved Δ*gcl* + BHAC strain. Two isoforms of isocitrate dehydrogenase (Icd and Idh) were upregulated, indicating increased flux through the TCA cycle, and eventually increased NADPH production. Together with the increased production of NAD(P) transhydrogenase, which can interconvert NADP(H) and NAD(H), this would ensure that the NAD(P)H requirements of the BHAC and biosynthetic reactions can be covered. In contrast, succinate-CoA ligase (SucCD) was downregulated, indicating decreased substrate-level phosphorylation in the evolved strain.

In amino acid metabolism, glutamate synthase levels (GltDB) were decreased, while dioxovalerate dehydrogenase (Dvd), which interconverts 2-OG and its semialdehyde, was slightly increased, as were the last two enzymes in lysine biosynthesis, DapF and LysA-I. The subunits of the glycine cleavage complex as well as serine dehydratase, which had already been upregulated in the parental Δ*gcl* + BHAC strain, were even further upregulated in the evolved strain. In contrast, the production of aspartate semialdehyde dehydrogenase (Asd) was decreased, indicating a decreased flux from this branching metabolite into lysine, methionine, threonine, and isoleucine biosynthesis. Finally, the levels of two enzymes in leucine biosynthesis (LeuB, LeuCD) was decreased as well, indicating a fine-tuning of amino acid biosynthesis in the evolved strain. In summary, analysis of the evolved Δ*gcl* + BHAC strain revealed pleiotropic effects that adapt the cellular networks of *P. putida* KT2440 for efficient integration of the BHAC.

## 4. Discussion

This work aimed at replacing the native glycerate pathway in *P. putida* KT2440 by a heterologous BHAC to establish a more efficient EG metabolism in this organism. To that end, we first implemented a linear BHA shunt in three different *E. coli* auxotrophic strains of increasing selection pressure. Having demonstrated that the BHA shunt was able to support complete biomass formation in *E. coli* Δ*gcl* after laboratory evolution, we subsequently transferred the full BHAC into a *P. putida* KT2440 Δ*gcl* strain. This allowed the strain to immediately grow on EG, notably at a similar growth rate and biomass yield as *P. putida* KT2440 E6.1, a strain that had recently been evolved to grow on EG via the glycerate pathway (Li et al., 2019). Laboratory evolution further improved growth of *P. putida* KT2440 Δ*gcl* + BHAC on EG. The final strain grows 35% faster than E6.1, with an increased maximum OD of 20%. Notably, the growth rate of the evolved Δ*gcl* + BHAC strain is not only improved compared to the previously evolved E6.1 strain (Li et al., 2019), but also to a *P. putida* strain that was engineered to overexpress the operons encoding for the glycerate pathway and glycolate oxidase (Franden et al., 2018).

Our work highlights that the transfer of complete heterologous metabolic modules combined with ALE is a powerful approach to overcome the inherent limitations of native metabolism and obtain strains with improved growth behavior. Genome sequencing and proteomics identified a multitude of changes as underlying factors of improved growth in the evolved *P. putida* KT2440 Δ*gcl* + BHAC strain. Notably, these changes are caused by genetic mutations, as well as plastic adaptations of the native genetic and metabolic network. Our studies with *E. coli* and *P. putida* KT2440 shed light on key changes that seem to be essential for an efficient integration of the BHAC. Three independently evolved *E. coli* Δ*gcl* strains showed point mutations in genes encoding succinate dehydrogenase (*sdhA*) or succinyl-CoA synthetase β-subunit (*sucC*), which enabled *E. coli* Δ*gcl* + BHAC to grow on glycine and glycolate with the BHA shunt. Notably, succinyl-CoA synthetase was also decreased in the evolved *P. putida* KT2440 Δ*gcl* + BHAC strain, likely as a result of altered regulation in central carbon metabolism.

Beyond succinate, there were other adaptations in central carbon metabolism, cellular redox regulation, the ETC, and amino acid biosynthesis that seemed important. Upon implementation of the pathway into *P. putida* KT2440, the native regulatory network immediately reacted and enzyme levels of the C3-C4 node shifted. This enabled the BHAC strain to shift from C3 assimilation (glycerate pathway) to C4 assimilation (BHAC) and made immediate E6.1-like growth of the engineered strain possible. Further adaptations required mutations of the regulatory network of *P. putida* KT2440, in particular, *lldR*, *regB* and a LysR-family regulator gene of unknown function, which showed pleiotropic effects on the metabolic network of *P. putida* KT2440. This multitude of changes to fully embed the BHAC with the genetic and metabolic network of the cell could most likely not have been realized by rational engineering, even though there is a broad variety of advanced genetic tools for *P. putida* (Martínez-García et al., 2017; Choi et al., 2020; Wirth et al., 2020; Patinios et al., 2021; Velázquez et al., 2021; Abdullah et al., 2022; Liu et al., 2022). In our view, the different changes in production of enzymes and respiratory complexes would have been nearly impossible to achieve by overexpressing genes or introducing point mutations. Therefore, the directed evolution approach will probably remain the method of choice to enable or improve complex phenotypes such as complete biomass synthesis via an engineered pathway.

The results of this study further advance the possibilities for microbial valorization of EG. This includes the upcycling of EG into biopolymers as reported recently in *Pseudomonas* (Kenny et al., 2008; Narancic et al., 2021; Tiso et al., 2021a), the microbial breakdown of PET in *Pseudomonas* (Narancic et al., 2021), as well as the engineering of biotechnologically and environmentally relevant *Pseudomonas* strains for improved growth. Notably, *P. umsongensis* GO16 was recently also evolved for better growth on EG (Tiso et al., 2021a). While this strain already shows a growth rate of 0.4 h^-1^, it is tempting to speculate that growth of *P. umsongensis* GO16 could be further improved by implementation of the BHAC in the future.

## Supporting information

Proteomics data 1 (parentBHAC vs E6-1)

Proteomics data 2 (evoBHAC vs parentBHAC)

## Data availability statement

Sequences from whole genome sequencing experiments have been deposited in the Sequence Read Archive (SRA) with the accession number PRJNA862906. All other relevant data are available in the article and the Supplementary Information.

## Author contributions

L.S.v.B., H.S.-M., M.T.C., F.S., and P.A.G.C. planned and performed experiments. T.G. performed mass spectrometry and data evaluation for proteomics. L.S.v.B., A.B.-E., S.N.L., and T.J.E. analyzed data and supervised the project. L.S.v.B. drafted the manuscript and prepared the final version together with H.S.-M., S.N.L., and T.J.E., with contributions from all other authors.

## Acknowledgements & Funding

We gratefully acknowledge Prof. Dr. Nick Wierckx (Forschungszentrum Jülich, Germany) for kindly providing us with *P. putida* KT2440 strains E6.1 and Δ*gcl* and the plasmid pBG35. We are grateful to Dr. José Vicente Gomes Filho (Philipps-University Marburg, Germany) for help with performing Illumina sequencing.

This study was funded by the Max-Planck-Society (Arren Bar-Even and Tobias J. Erb), the European Union Horizon 2020 research and innovation programme (Grant Agreement 862087, ‘Gain4Crops’), and the German Research Foundation (SFB987 ‘Microbial diversity in environmental signal response’).

