## Supplementary Information for "Implementation of the β-hydroxyaspartate cycle increases growth performance of *Pseudomonas putida* on the PET monomer ethylene glycol"

*[1]Department of Biochemistry & Synthetic Metabolism, Max Planck Institute for Terrestrial Microbiology, Karl-von-Frisch-Str. 10, 35043 Marburg, Germany; [2]Institute of Biology Leiden, Leiden University, Sylviusweg 72, 2333 BE Leiden, The Netherlands; [3]Max Planck Institute of Molecular Plant Physiology, Am Mühlenberg 1, 14476 Potsdam-Golm, Germany; [4]Facility for Mass Spectrometry and Proteomics, Max Planck Institute for Terrestrial Microbiology, Karl-von-Frisch-Str. 10, 35043 Marburg, Germany; [5]Department of Biochemistry, Charité Universitätsmedizin, Charitéplatz 1, 10117 Berlin, Germany; [6]LOEWE-Center for Synthetic Microbiology, Philipps-University Marburg, Karl-von-Frisch-Str. 8, 35043 Marburg, Germany.*

*# present address: School of Biomolecular and Biomedical Sciences and BiOrbic SFI Bioeconomy Research Centre, University College Dublin, Belfield, Ireland*

*† these authors contributed equally*

**Table of Contents Supplementary Information:**

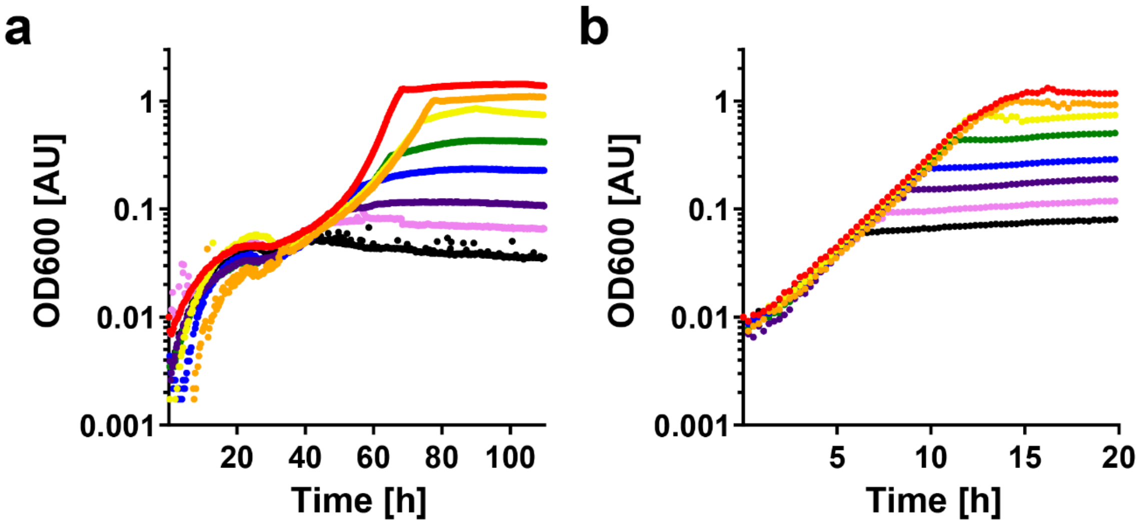

**SI Figure 1: Supplementation of the aspartate auxotrophic strain with aspartate and tyrosine. a**, Growth of the aspartate auxotrophic strain on M9 minimal medium with 20 mM glycerol and different concentrations of aspartate (from red curve to black curve: 5, 2.9, 1.6, 0.9, 0.5, 0.3, 0.17, 0.1 mM). **b**, Growth of the aspartate auxotrophic strain on M9 minimal medium with 20 mM glycerol, 1 mM tyrosine, and different concentrations of aspartate (see above).

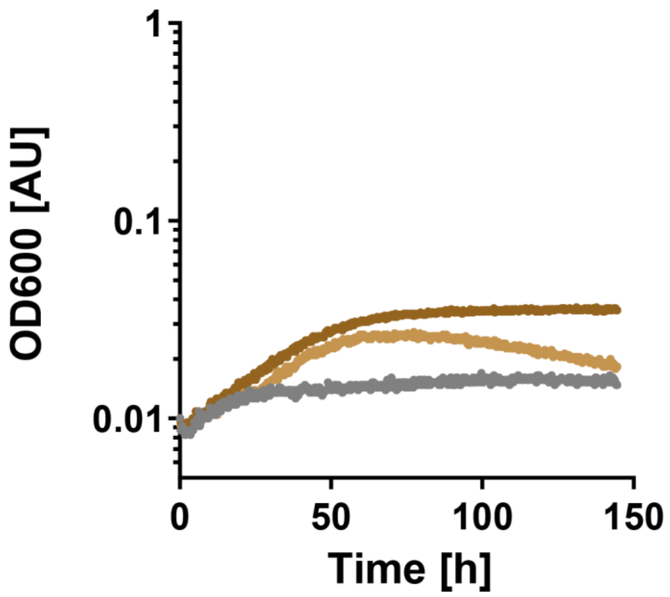

**SI Figure 2: Growth controls of the oxaloacetate auxotrophic strain + BHA shunt.** Growth of the oxaloacetate auxotrophic strain + BHA shunt on M9 minimal medium with 20 mM glycerol and 5 mM glyoxylate + 5 mM glycine (light brown), 5 mM glycolate + 5 mM glycine (brown), or 5 mM 2-OG (gray).

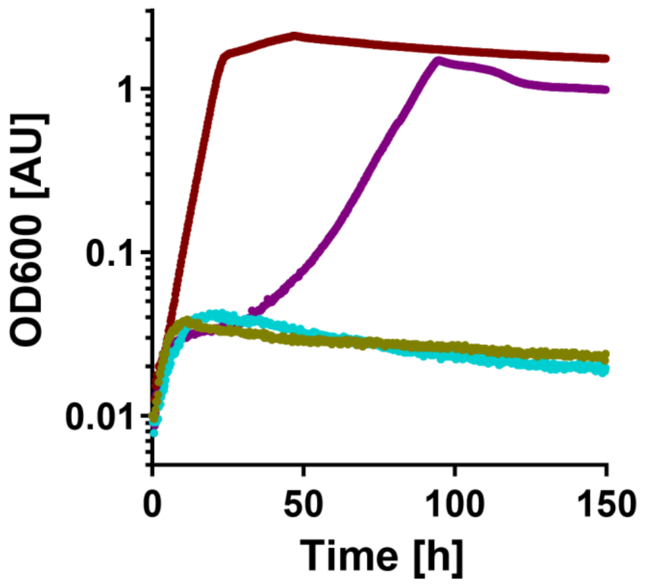

**SI Figure 3: Growth controls of the evolved isolate of Δ*gcl* + BHA shunt.** Growth of the evolved isolate of Δ*gcl* + BHA shunt on M9 minimal medium with 80 mM glycolate (turquoise), 10 mM glycine (olive), 80 mM glycolate + 10 mM glycine (purple), and 20 mM glycerol (maroon).

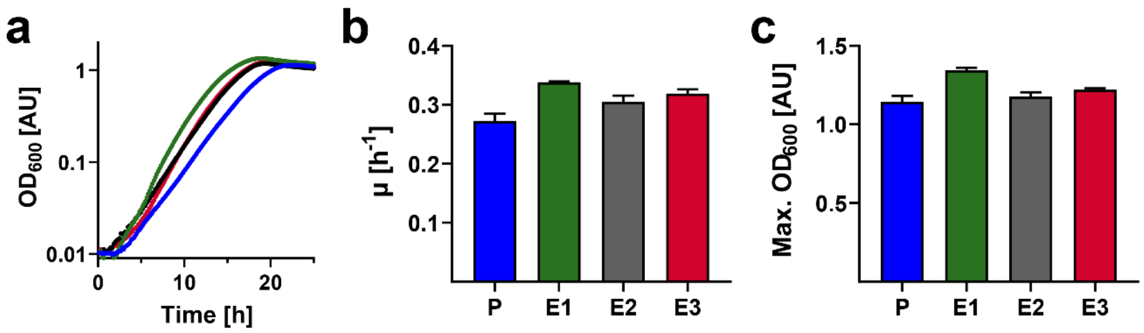

**SI Figure 4: Growth of evolved *P. putida* KT2440 Δ*gcl* + BHAC strains. a**, Growth curves of Δ*gcl* + BHAC (blue), and its evolved derivatives (green, gray, red) grown in the presence of 20 mM EG. Shown are representative growth curves from n = 6 independent cultures, with errors < 5%. **b**, Growth rates (µ) and **c**, Max. $OD_{600}$ of Δ*gcl* + BHAC (parental strain and three evolved isolates) grown in the presence of 20 mM EG. Data are the mean ± s.d. of n = 6 independent cultures. Strain E1 exhibited the best growth performance and was therefore selected for in-depth characterization.

### Supplementary Table 1 | Strains and plasmids used in this study

| strain or plasmid | genotype or relevant features[a] | source or reference |
|---|---|---|
| *E. coli* DH5α | *supE44*, *ΔlacU169* (*Φ80lacZDM15*), *hsdR17*, *recA1*, *endA1*, *gyrA96*, *thi*-1, *relA1* | Thermo Fisher Scientific, Waltham, USA |
| *E. coli* BL21 AI | *ompT*, *gal*, *dcm*, *lon*, *hsdS*$_B$($r_B^- m_B^-$), [*malB*$^+$]$_{K\text{-}12}$($\lambda^S$), *araB*::*T7RNAP-tetA* | Thermo Fisher Scientific, Waltham, USA |
| *E. coli* SIJ488 | integrated λ-red recombinase and flippase | Jensen et al. (2015) |
| *E. coli* oxaloacetate auxotroph | Δ*aspA* Δ*ppc* Δ*pck* Δ*mdh* Δ*mqo* | this work |
| *E. coli* aspartate auxotroph | Δ*aspC* Δ*tyrB* | this work |
| *E. coli* Δ*gcl* | Δ*gcl* | this work |
| *E. coli* Δ*gcl* pZ-ASS-BhcCBD mutant 1 | Δ*gcl* pZ-ASS-BhcBCD evolved to grow on glycolate + glycine; carries mutations in *sdhA, kgtP, yhiM, entF* | this work |
| *E. coli* Δ*gcl* pZ-ASS-BhcCBD mutant 2 | Δ*gcl* pZ-ASS-BhcBCD evolved to grow on glycolate + glycine; carries mutation in *sucC* | this work |
| *E. coli* Δ*gcl* pZ-ASS-BhcCBD mutant 3 | Δ*gcl* pZ-ASS-BhcBCD evolved to grow on glycolate + glycine; carries mutation in *sucC* | this work |
| *E. coli* JW4014 | Keio Δ*tyrB* | Baba et al. (2006) |
| *E. coli* JW0495 | Keio Δ*gcl* | Baba et al. (2006) |
| *P. putida* KT2440 E6.1 | *P. putida* KT2440 evolved to grow on ethylene glycol; carries mutations in *gclR*, PP_2046, PP_2662 | Li et al. (2019) |
| *P. putida* KT2440 Δ*gcl* | Δ*gcl* | Wierckx lab, FZ Jülich, Jülich, Germany |
| *P. putida* KT2440 Δ*gcl* pBG35-BhcABCD mutant | Δ*gcl* pBG35-BhcABCD evolved to grow more efficiently on ethylene glycol; carries mutations in *regB*, PP_1861, *flhB*, *lldR* | this work |
| pNivC | Cloning vector; $Amp^R$ | Zelcbuch et al. (2013) |
| pZ-ASS | Overexpression plasmid with p15A origin and constitutive strong promoter; $Str^R$ | Wenk et al. (2018) |
| pZ-ASS-BhcCBD | pZ-ASS backbone for overexpression of β-hydroxyaspartate aldolase, β-hydroxyaspartate dehydratase and iminosuccinate reductase from *P. denitrificans*; $Str^R$ | this work |
| pZ-ASS-BhcCB | pZ-ASS backbone for overexpression of β-hydroxyaspartate aldolase and β-hydroxyaspartate dehydratase from *P. denitrificans;* $Str^R$ | this work |
| pTE104 | Broad-host range expression vector, *P*coxB promoter, IncP' and pMB1 origins; $Tc^R$ | Schada von Borzyskowski et al. (2015) |
| pBG35 | mini-Tn7 transposon, translational coupler, synthetic promoter 14b, R6K origin; $Gm^R$, $Km^R$ | Zobel et al. (2015) |
| pTE104-BhcABCD | pTE104 backbone for overexpression of aspartate-glyoxylate aminotransferase, β-hydroxyaspartate aldolase, β-hydroxyaspartate dehydratase and iminosuccinate reductase from *P. denitrificans*; $Tc^R$ | this work |
| pBG35-BhcABCD | pTE104 backbone (*P*coxB replaced with *P*BG35) for overexpression of aspartate-glyoxylate aminotransferase, β-hydroxyaspartate aldolase, β-hydroxyaspartate dehydratase and iminosuccinate reductase from *P. denitrificans*; $Tc^R$ | this work |

[a] $Amp^R$, ampicillin resistance; $Str^R$, streptomycin resistance; $Tc^R$, tetracycline resistance; $Gm^R$, gentamicin resistance; $Km^R$, kanamycin resistance

**Supplementary Table 2 | Primers used in this study.** 'KO' primers were used to amplify the CapR knockout cassette from pKD3 with 50 bp gene-specific upstream and downstream sequences. 'KO-Ver'-primers (knockout-verification) were used to verify gene replacement by chloramphenicol or kanamycin resistance cassettes and cassette removal by flippase. Internal primers ('int') were used to verify successful removal of the gene from the genome.

| target | name | sequence |
|---|---|---|
| Eco_*aspA* | aspA_KO_fwd | GCCTTTTTTATTTGTACTACCCTGTACGATTACTGTTCGCTTTCATCAGTGTGTAGGCTGGAGCTGCTTC |
| Eco_*aspA* | aspA_KO_rvs | CTGTGTGTTTAAAGCAAATCATTGGCAGCTTGAAAAAGAAGGTTCACATGCATATGAATATCCTCCTTAG |
| Eco_*aspA* | aspA_KO_Ver_fwd | TCCGCCTGCAAAACCAATACCT |
| Eco_*aspA* | aspA_KO_Ver_rvs | CTCGGGTATTCGGTCGATGCAG |
| Eco_*aspA* | aspA_int_fwd | TCCGCTTCAGTCAACAGACC |
| Eco_*aspA* | aspA_int_rvs | GGAAGTTCCAGCTGATGCCT |
| Eco_*aspC* | aspC_KO_fwd | TTTTCAGCGGGCTTCATTGTTTTTAATGCTTACAGCACTGCCACAATCGCGTGTAGGCTGGAGCTGCTTC |
| Eco_*aspC* | aspC_KO_rvs | TACCCTGATAGCGGACTTCCCTTCTGTAACCATAATGGAACCTCGTCATGCATATGAATATCCTCCTTAG |
| Eco_*aspC* | aspC_KO_Ver_fwd | GCCTGCATAATCCCTTCCTGCA |
| Eco_*aspC* | aspC_KO_Ver_rvs | GTCTTGCAAAAACAGCCTGCGT |
| Eco_*ppc* | ppc_KO_fwd | GAAGGATACAGGGCTATCAAACGATAAGATGGGGTGTCTGGGGTAATATGAATTAACCCTCACTAAAGGGCG |
| Eco_*ppc* | ppc_KO_rvs | AAAGCACGAGGGTTTGCAGAAGAGGAAGATTAGCCGGTATTACGCATACCTAATACGACTCACTATAGGGCTC |
| Eco_*ppc* | ppc_KO_V_fwd | ACGAGGGTGTTAGAACAGAAGT |
| Eco_*ppc* | ppc_KO_V_rvs | CAAAGCCCGAGCATATTCGC |
| Eco_*ppc* | ppc_int_fwd | CGGTATTACGCATACCTGCCG |
| Eco_*ppc* | ppc_int_rev | TCAGGGCAAACAGATGGTGAT |
| Eco_*pck* | pck_KO_F | CTGGATAGATATTCTCCAGCTTCAAATCATTACAGTTTCGGACCAGCCGCAATTAACCCTCACTAAAGGGCG |
| Eco_*pck* | pck_KO_R | AAAGACTTTACTATTCAGGCAATACATATTGGCTAAGGAGCAGTGAAATGTAATACGACTCACTATAGGGCTC |
| Eco_*pck* | pck_KO_V_F | ATCTATGAGCCTTGTCGCGG |
| Eco_*pck* | pck_KO_V_R | GCAGGGCACGACAAAAGAAG |
| Eco_*pck* | pck_int_rev | AAGCATCAGCAGTCAGGAAGA |
| Eco_*pck* | pck_int_fwd | CCAAGCTACGACCTGCTGTAT |
| Eco_*mdh* | mdh_KO_fwd | GCTCCGGTTTTTTTATTATCCGCTAATCAATTACTTATTAACGAACTCTTCGTGTAGGCTGGAGCTGCTTC |
| Eco_*mdh* | mdh_KO_rvs | CAGCGGAGCAACATATCTTAGTTTATCAATATAATAAGGAGTTTAGGATGGGAATTAGCCATGGTCCATATG |
| Eco_*mdh* | mdh_KO_V_fwd | CGTGGTTAATGAAGTGTCACC |
| Eco_*mdh* | mdh_KO_V_rvs | TTGTCACCACCTGTTGGAATGTT |
| Eco_*mdh* | mdh_int_fwd | GTGCACGAACCAGAGACAGA |
| Eco_*mdh* | mdh_int_rvs | GGGTATGGATCGTTCCGACC |
| Eco_*mqo* | mqo_KO_fwd | CGGCATACCATGCCGGATGTGGCGTATCATTACAACGCAATATCCGCCACGTGTAGGCTGGAGCTGCTTC |
| Eco_*mqo* | mqo_KO_rvs | GCAAATCTCACCTGCAATAAAGCGACTAAAAGTAAGGCATTAACAAGATGCATATGAATATCCTCCTTAG |
| Eco_*mqo* | mqo_KO_V_fwd | GCCGCATCCAACATCTAACG |
| Eco_*mqo* | mqo_KO_V_rvs | CGGACTGCTGCCGTCA |
| Eco_*mqo* | mqo_int_fwd | CGGTTTCAACTGCGGTTTCG |
| Eco_*mqo* | mqo_int_rvs | GGAACTGAACTACACCCCGC |
| Eco_*tyrB* | tyrB_KO_Ver_fwd | ACATCCACTCGATCTTCGCC |

| | | |
|---|---|---|
| Eco_*tyrB* | tyrB_KO_Ver_rvs | TCACGTAGAACGATGGCATCA |
| Eco_*tyrB* | tyrB_int_fwd | TCAGCGATCCTACCTGGGAA |
| Eco_*tyrB* | tyrB_int_rvs | TTGCGTTCTGGCATCTCTGT |
| Eco_*gcl* | gcl_KO_Ver_fwd | CCGGGCCTTCATCAAGACTG |
| Eco_*gcl* | gcl_KO_Ver_rvs | CGCCGGTAAATTGTGCAGCG |
| Eco_*gcl* | gcl_int fwd | GTCATGTGGAAGGTGCTTCGC |
| Eco_*gcl* | gcl_int_rev | AGCATTTGTGCCGCAGCGATT |
| Sequencing | pZ-ASS-seq-fwd | GCATTTATCAGGGTTATTGTCTCATG |
| Sequencing | Cap-seq-rvs | CTGAACGGTCTGGTTATAGG |
| Pden_*bhcAB* | bhcAB_fw | GCGGGGACATAAGCTTAGATCTTGAACAAGCGCAGGAGGAATTGCC |
| Pden_*bhcAB* | bhcAB_rv | CCGTTTTCGCGTTCATGGGGCTCTCCTCAATTCTGGTTC |
| Pden_*bhcCD* | bhcCD_fw | TGAGGAGAGCCCCATGAACGCGAAAACGGATTTC |
| Pden_*bhcCD* | bhcCD_rv | GCCAGTGAATTAGGTACCTGCAGGAGATCCCGCAGACGAAAAGGTC |
| *P*BG35 | *P*BG35_Hyb_fw | CTAGTTAATTAATTTATTTGACATGCGTGATGTTTAGAATTATAATTTGGGGACCTAGG |
| *P*BG35 | *P*BG35_Hyb_rv | AGCTCCTAGGTCCCCAAATTATAATTCTAAACATCACGCATGTCAAATAAATTAATTAA |
| Sequencing | Brick_fw_seq | CCTATGGAAAAACGCCAGCAACGC |
| Sequencing | Brick_rv_seq | TCGCCATTCAGGCTGCGCAACTGTTGG |

### **Sequences of genes codon-optimized for *E. coli***

Aspartate-glyoxylate aminotransferase (BhcA, Uniprot A1B8Z3) from *Paracoccus denitrificans*

ATGCATCATCACCATCACCACACCTCTCAGAACCCGATCTTCATCCCGGGTCCGACCAACATCCCGGAAGAAAT
GCGTAAAGCTGTTGACATGCCGACCATCGACCACCGTTCTCCGGTTTTCGGTCGTATGCTGCACCCGGCTCTGG
AAGGTGTTAAAAAAGTTCTGAAAACCACCCAGGCTCAGGTTTTCCTGTTCCCGTCTACCGGTACCGGTGGTTGG
GAAACCGCTATCACCAACACCCTGTCTCCGGGTGACAAAGTTCTGGCTGCTCGTAACGGTATGTTCTCTCACCG
TTGGATCGACATGTGCCAGCGTCACGGTCTGGACGTTACCTTCGTTGAAACCCCGTGGGGTGAAGGTGTTCCG
GCTGACCGTTTCGAAGAAATCCTGACCGCTGACAAAGGTCACGAAATCCGTGTTGTTCTGGCTACCCACAACG
AAACCGCTACCGGTGTTAAATCTGACATCGCTGCTGTTCGTCGTGCTCTGGACGCTGCTAAACACCCGGCTCTG
CTGTTCGTTGACGGTGTTTCTTCTATCGGTTCTATGGACTTCCGTATGGACGAATGGGGTGTTGACATCGCTGT
TACCGGTTCTCAGAAAGGTTTCATGCTGCCGCCGGGTCTGGCTATCGTTGGTTTCTCTCCGAAAGCTATGGAAG
CTGTTGAAACCGCTCGTCTGCCGCGTACCTTCTTCGACATCCGTGACATGGCTACCGGTTACGCTCGTAACGGT
TACCCGTACACCCCGCCGGTTGGTCTGATCAACGGTCTGAACGCTTCTTGCGAACGTATCCTGGCTGAAGGTCT
GGAAAACGTTTTCGCTCGTCACCACCGTATCGCTTCTGGTGTTCGTGCTGCTGTTGACGCTTGGGGTCTGAAAC
TGTGCGCTGTTCGTCCGGAACTGTACTCTGACTCTGTTTCTGCTATCCGTGTTCCGGAAGGTTTCGACGCTAACC
TGATCGTTTCTCACGCTCTGGAAACCTACGACATGGCTTTCGGTACCGGTCTGGGTCAGGTTGCTGGTAAAGTT
TTCCGTATCGGTCACCTGGGTTCTCTGACCGACGCTATGGCTCTGTCTGGTATCGCTACCGCTGAAATGGTTAT
GGCTGACCTGGGTCTGCCGATCCAGCTGGGTTCTGGTGTTGCTGCTGCTCAGGAACACTACCGTCAGACCACC
GCTGCTGCTCAGAAAAAAGCTGCTTAATCTAGAGCTAGC

β-hydroxyaspartate aldolase (BhcC, Uniprot A1B8Z1) from *Paracoccus denitrificans*

ATGCATCATCACCATCACCACAACGCTAAAACCGACTTCTCTGGTTACGAAGTTGGTTACGACATCCCGGCTCT
GCCGGGTATGGACGAATCTGAAATCCAGACCCCGTGCCTGATCCTGGACCTGGACGCTCTGGAACGTAACATC
CGTAAAATGGGTGACTACGCTAAAGCTCACGGTATGCGTCACCGTTCTCACGGTAAAATGCACAAATCTGTTG
ACGTTCAGAAACTGCAAGAATCTCTGGGTGGTTCTGTTGGTGTTTGCTGCCAGAAAGTTTCTGAAGCTGAAGCT
TTCGCTCGTGGTGGTATCAAAGACGTTCTGGTTACCAACGAAGTTCGTGAACCGGCTAAAATCGACCGTCTGG
CTCGTCTGCCGAAAACCGGTGCTACCGTTACCGTTTGCGTTGACGACGTTCAGAACATCGCTGACCTGTCTGCT
GCTGCTCAGAAACACGGTACCGAACTGGGTATCTTCGTTGAAATCGACTGCGGTGCTGGTCGTTGCGGTGTTA
CCACCAAAGAAGCTGTTGTTGAAATCGCTAAAGCTGCTGCTGCTGCTCCGAACCTGACCTTCAAAGGTATCCAG
GCTTACCAGGGTGCTATGCAGCACATGGACTCTTTCGAAGACCGTAAAGCTAAACTGGACGCTGCTATCGCTC
AGGTTAAAGAAGCTGTTGACGCTCTGGAAGCTGAAGGTCTGGCTCCGGAATTTGTTTCTGGTGGTGGTACCGG
TTCTTACTACTTCGAATCTAACTCTGGTATCTACAACGAACTGCAATGCGGTTCTTACGCTTTCATGGACGCTGA
CTACGGTCGTATCCACGACGCTGAAGGTAAACGTATCGACCAGGGTGAATGGGAAAACGCTCTGTTCATCCTG
ACCTCTGTTATGTCTCACGCTAAACCGCACCTGGCTGTTGTTGACGCTGGTCTGAAAGCTCAGTCTGTTGACTCT
GGTCTGCCGTTCGTTTACGGTCGTGACGACGTTAAATACATCAAATGCTCTGACGAACACGGTGTTGTTGAAG
ACAAAGACGGTGTTCTGAAAGTTAACGACAAACTGCGTCTGGTTCCGGGTCACTGCGACCCGACCTGCAACGT
TCACGACTGGTACGTTGGTGTTCGTAACGGTAAAGTTGAAACCGTTTGGCCGGTTTCTGCTCGTGGTAAAGGT
TACTAATCTAGAGCTAGC

β-hydroxyaspartate dehydratase (BhcB, Uniprot A1B8Z2) from *Paracoccus denitrificans*

ATGCATCATCACCATCACCACTACATCCCGACCTACGAAGACATGCTGGCTGCTCACGAACGTATCAAACCGCA
CATCCGTCGTACCCCGATCCGTACCTCTGACTACCTGAACGAACTGACCGGTGCTCAGCTGTTCTTCAAATGCG
AAAACTTCCAGGAACCGGGTGCTTTCAAAGTTCGTGGTGCTACCAACGCTGTTTTCGGTCTGGACGACGCTCA
GGCTGCTAAAGGTGTTGCTACCCACTCTTCTGGTAACCACGCTTCTTGCCTGTCTTACGCTGCTATGCTGCGTGG
TATCCCGTGCAACGTTGTTATGCCGCGTACCGCTCCGCAGGCTAAAAAAGACACCGTTCGTCGTTACGGTGGT

GTTATCACCGAATGCGAACCGTCTACCTCTTCTCGTGAAGAAACCTTCGCTAAAGTTCAGGCTGAAACCGGTGG
TGACTTCGTTCACCCGTACAACGACCCGCGTGTTATCGCTGGTCAGGGTACCTGCGCTAAAGAACTGGTTGAAC
AGGTTGACGGTCTGGACGCTGTTGTTGCTCCGATCGGTGGTGGTGGTATGATCTCTGGTACCTGCCTGACCCT
GTCTACCCTGGCTCCGGAAACCCGTGTTATCGCTGCTGAACCGGAACAGGCTGACGACGCTTACCGTTCTTTCA
AAGCTGGTTACATCATCGCTGACGACGCTCCGAAAACCGTTGCTGACGGTCTGCTGGTTCCGCTGAAAGACCT
GACCTGGCACTTCGTTAAAAACCACGTTTCTGAAATCTACACCGCTTCTGACGCTGAAATCGTTGACGCTATGA
AACTGATCTGGAAACACCTGCGTATCGTTATGGAACCGTCTTCTGCTGTTCCGCTGGCTACCATCCTGAAAAAC
CCGGAAGCTTTCGCTGGTAAACGTGTTGGTGTTATCGTTACCGGTGGTAACGTTGACCTGGACAAACTGCCGT
GGAACTAATCTAGAGCTAGCG

Iminosuccinate reductase (BhcD, Uniprot A1B8Z0) from *Paracoccus denitrificans*

ATGCATCATCACCATCACCACCTGGTTGTTGCTGAAAAAGAAATCGCTGGTCTGATGACCCCGGAAGCTGCTTT
CGAAGCTATCGAAGCTGTTTTCGCTTCTATGGCTCGTCGTAAAGCTTACAACTTCCCGGTTGTTCGTGAAGCTAT
CGGTCACGAAGACGCTCTGTACGGTTTCAAAGGTGGTTTCGACGCTTCTGCTCTGGTTCTGGGTCTGAAAGCT
GGTGGTTACTGGCCGAACAACCAGAAACACAACCTGATCAACCACCAGTCTACCGTTTTCCTGTTCGACCCGGA
CACCGGTCGTGTTTCTGCTGCTGTTGGTGGTAACCTGCTGACCGCTCTGCGTACCGCTGCTGCTTCTGCTGTTTC
TATCAAATACCTGGCTCCGAAAGGTGCTAAAGTTCTGGGTATGATCGGTGCTGGTCACCAGTCTGCTTTCCAGA
TGCGTGCTGCTGCTAACGTTCACCGTTTCGAAAAAGTTATCGGTTGGAACCCGCACCCGGAAATGCTGTCTCGT
CTGGCTGACACCGCTGCTGAACTGGGTCTGCCGTTCGAAGCTGTTGAACTGGACCGTCTGGGTGCTGAAGCTG
ACGTTATCGTTTCTATCACCTCTTCTTTCTCTCCGCTGCTGATGAACGAACACGTTAAAGGTCCGACCCACATCG
CTGCTATGGGTACCGACACCAAAGGTAAACAGGAACTGGACCCGGCTCTGGTTGCTCGTGCTCGTATCTTCAC
CGACGAAGTTGCTCAGTCTGTTTCTATCGGTGAATGCCAGCACGCTATCGCTGCTGGTCTGATCCGTGAAGACC
AGGTTGGTGAACTGGGTGCTGTTGTTGCTGGTGACGACCCGGGTCGTGGTGACGCTGAAGTTACCATCTTCGA
CGGTACCGGTGTTGGTCTGCAAGACCTGGCTGTTGCTCAGGCTGTTGTTGAACTGGCTAAACACAAAGGTGTT
GCTCAGGAAGTTGAAATCTAATCTAGAGCTAGC
